# Metabolic Crossroads: AMDHD2 Couples the Hexosamine Biosynthetic Pathway to Acetyl-CoA Homeostasis in Pluripotent Stem Cells

**DOI:** 10.1101/2025.08.20.671247

**Authors:** Virginia Kroef, Lea Hund, Yasmin Barenco Abbas, Norito Tamura, Jenny Fröhlich, Ann-Kathrin Huber, Syed Musa Ali, Lena Wischhof, Hannah Scheiblich, Patrick Giavalisco, Martin S. Denzel, Peter Tessarz, Ina Huppertz

## Abstract

Pluripotent stem cells (SCs) rely on metabolic rewiring to regulate self-renewal and differentiation. The hexosamine biosynthetic pathway (HBP) integrates multiple metabolic inputs to produce uridine diphosphate-N-acetylglucosamine (UDP-GlcNAc), a central substrate for protein and lipid glycoconjugates. While UDP-GlcNAc levels influence pluripotency, the broader metabolic roles of the HBP in stem cell maintenance and differentiation remain largely unexplored. Using proteomic and metabolomic analyses, we uncovered extensive enzymatic remodelling of the HBP during early mouse embryonic stem cell (mESC) differentiation, including a significant decrease in N-acetylglucosamine-6-phosphate (GlcNAc6P) levels, despite stable UDP-GlcNAc levels. Isotope labelling experiments revealed that pluripotent stem cells maintain UDP-GlcNAc biosynthesis through a combination of glucose-driven *de novo* synthesis and the N-Acetylglucosamine (GlcNAc) salvage pathway, both of which are downregulated during differentiation. Functionally, we identified GlcNAc6P—a point of convergence of the *de novo* synthesis and the salvage pathways—as a pivotal metabolite of stem cell fate. Preventing GlcNAc6P’s deacetylation by deleting or depleting AMDHD2 impaired mESC differentiation. Mechanistically, we demonstrated that GlcNAc6P catabolism is essential for sustaining cytosolic acetate and in turn acetyl-CoA levels, which are critical for lipid synthesis and histone modifications required for pluripotency exit. These findings establish AMDHD2 as a key supplier of cellular acetate and reveal an essential metabolic link between HBP activity and acetyl-CoA homeostasis. By expanding the role of the HBP beyond UDP-GlcNAc production, this study reveals an unrecognised metabolic vulnerability that can be exploited to manipulate stem cell identity.

## Introduction

Pluripotent stem cells (SCs) are characterized by a unique metabolic profile that requires rewiring during the exit from pluripotency and early differentiation to adapt to their novel cellular functional properties^1–3^. While SCs mainly utilize aerobic glycolysis to meet their high energy demands, also known as the Warburg effect, differentiated cells preferentially rely on oxidative phosphorylation (OXPHOS) for energy production^1, 2, 4^. Since SC metabolism is tightly linked to cell identity and fate decisions, cellular metabolism requires strict regulation. For example, core pluripotency and transcription factors such as OCT4, SOX2 and NANOG are known to regulate glucose metabolism by transcriptionally upregulating the expression of glycolytic enzymes, highlighting the link between metabolism and pluripotency^4–6^. Furthermore, reduced glycolytic flux during differentiation is accompanied by a decrease in acetyl-CoA levels, which are essential for histone acetylation, leading to a transcriptional reprogramming that supports metabolic rewiring of SCs^7^. Therefore, maintaining elevated acetyl-CoA levels in pluripotent cells is essential, while persistently high levels can disrupt differentiation^7^. Tightly regulated metabolic networks have evolved to preserve cellular acetyl-CoA homeostasis. For example, loss of ATP-citrate synthase (ACLY), which converts citrate into cytosolic acetyl-CoA, leads to increased Acyl-CoA Synthetase Short Chain Family Member 2 (ACSS2) expression in various cancer cell lines and MEFs, providing an alternative route to generate acetyl-CoA by utilizing acetate^8–11^.

The hexosamine biosynthetic pathway (HBP) is a metabolic pathway that consumes approximately 2–5% of cellular glucose to generate substrates for various post-translational modifications, such as O-GlcNAcylation, N-glycosylation and O-glycosylation, and thus plays an essential role in metabolic adaptations and cellular homeostasis, particularly in rapidly dividing cells^12–14^.

In the first and rate-limiting step of the HBP, the glycolysis intermediate fructose-6-phosphate (Frc6P) together with L-glutamine is converted to D-glucosamine-6-phosphate (GlcN6P) by the enzymes glutamine-fructose-6-phosphate amidotransferase1/2 (GFAT1/2 or GFPT1/2)^13^. GlcN6P is then used as a substrate for acetylation by the enzyme glucosamine-phosphate N-acetyltransferase (GNPNAT1), which produces N-acetylglucosamine-6-phosphate (GlcNAc6P) by transferring the acetyl group from acetyl-coenzyme A (acetyl-CoA)^15^. These first two steps of the HBP can be reversed by the action of the enzymes N-acetylglucosamine deacetylase 2 (AMDHD2 or amidohydrolase domain containing 2) and glucosamine-6-P deaminase 1/2 (GNPDA1/2), which remove the acetyl- and amide groups, respectively, forming the counteractors of GNPNAT1 and GFPT1/2^16–18^. Subsequently, the isomerization of GlcNAc6P to GlcNAc-1-phosphate (GlcNAc1P) is mediated by GlcNAc phosphomutase 3 (PGM3) and, in a final step, UDP is added by UDP-N-acetylglucosamine pyrophosphorylase 1 (UAP1) to synthesize the end product of the HBP, UDP-GlcNAc^19, 20^. Since the *de novo* synthesis of UDP-GlcNAc requires the consumption of intermediates of major metabolic pathways (Frc6P, L-glutamine, UTP and acetyl-CoA), we hypothesized that the HBP may act as a nutrient sensor coordinating important downstream processes^12, 21^. In cancer cells it has been shown that UDP-GlcNAc production depends not only on *de novo* synthesis, but also on intracellular GlcNAc, which can be derived from several sources, including the removal of O-GlcNAc modifications, degradation of glycoconjugates, and lysosomal breakdown of extracellular glycoproteins following micropinocytosis^22^. Subsequently, GlcNAc is converted through phosphorylation by N-acetyl-D-glucosamine kinase (NagK) to produce GlcNAc6P, which enters the HBP and bypasses the rate-limiting step of *de novo* synthesis, a process known as the salvage pathway^22^.

The HBP has been previously described to regulate cell identity and SC maintenance, mainly through its end product UDP-GlcNAc. Specifically, UDP-GlcNAc-mediated O-GlcNAcylation was shown to positively regulate the function of the key pluripotency factors OCT4 and SOX2 in pluripotent cells^23–26^. In addition, several studies have shown that transcriptional and epigenetic profiles in ESCs can be regulated by UDP-GlcNAc, either through O-GlcNAcylation of the DNA demethylation enzyme ten-eleven translocation 1 (TET1), the Polycomb repressive complex member EZH2, or the core histones^27–30^. Furthermore, UDP-GlcNAc is essential for the synthesis of constituent macromolecules of the extracellular matrix (ECM), which plays a regulatory role in SC fate^31^. In our previous work, we demonstrated a switch in the HBP enzyme expression levels when comparing pluripotent mESCs and terminally differentiated cells. While GFPT2 plays an important role in regulating UDP-GlcNAc levels in pluripotent cells, higher levels of the isozyme GFPT1 but no detectable GFPT2 levels were found in terminally differentiated cells^32^. Thus, HBP is an integral component of SC maintenance and subject to strictly coordinated regulation during differentiation.

In this study, we set out to understand how the HBP is rewired during early mESC differentiation, how this affects other related pathways and whether the HBP may regulate SC fate decisions independently of UDP-GlcNAc. To this end, we performed metabolic and proteomic analyses of mESCs during spontaneous differentiation and identified a drastic rearrangement of major pathways such as glycolysis and the TCA cycle, and more prominently, the HBP. We discovered that, in addition to *de novo* synthesis, GlcNAc recycling through the salvage pathway plays a critical role in the generation of UDP-GlcNAc, with both processes decreasing during differentiation. In addition, we showed that GlcNAc6P, as the convergence point of both pathways, is a key molecule for SC differentiation and that altering its deacetylation by deleting or depleting the corresponding enzyme AMDHD2, was sufficient to disrupt mESC differentiation.

We discovered that the deacetylation of GlcNAc6P by AMDHD2 is a source of acetate and contributes to the levels of acetyl-CoA downstream in pluripotent cells. Furthermore, we demonstrated that AMDHD2 KO cells attempt to compensate for the loss of acetate production by alternative mechanisms. These include decreased fatty acid synthase (FASN) and increased ACSS2 levels, which are comparable to the response described for ACLY loss^8–10^. Importantly, supplementing acetate in AMDHD-depleted cells restored productive stem cell differentiation. Furthermore, inhibiting the conversion of acetate to acetyl-CoA in wild-type (WT) cells using an ACSS2 inhibitor induced a compromised differentiation phenotype similar to that observed in AMDHD2 KO cells. Finally, we could show that acetate produced by AMDHD2 can be used in lipid biosynthesis and that loss of AMDHD2-derived acetate leads to altered histone acetylation and methylation patterns in the promoter regions of pluripotency and differentiation genes.

Together, these findings identify AMDHD2—and by extension, the HBP— as a novel regulatory axis that limits acetate levels in pluripotent cells. This mechanism not only deepens our understanding of the metabolic control of stem cell fate but also suggests potential avenues for manipulating stem cell differentiation in regenerative medicine.

## Results

### AN3-12 mouse embryonic stem cells display metabolic rewiring upon the removal of the leukaemia inhibitory factor

During the exit from pluripotency, mESCs undergo a metabolic shift which is accompanied by a rewiring of the major metabolic pathways, like glycolysis, the pentose phosphate pathway (PPP), the tricarboxylic acid (TCA) cycle, and OXPHOS, to adapt to their new cellular functions and properties^33, 34^. Since the HBP integrates metabolites from all these important pathways to generate UDP-GlcNAc, detailed information about how the pathway is rewired during differentiation and how this can impact stem cell fate, needs to be examined. Based on our previous work, we know that the HBP is part of a complex regulatory network, with an isozyme switch occurring during the exit of pluripotency (GFPT2 to GFPT1), limiting basal UDP-GlcNAc levels by increased feedback inhibition^32^. We used pluripotent AN3-12 mESCs cultivated in medium containing the leukaemia inhibitory factor (LIF) (referred to as D0) and induced differentiation by removing LIF from the media for one, three, five, or seven days (D1, D3, D5, D7) to systematically investigate how the HBP enzymes are altered as mESCs undergo differentiation (Figure 1A). The withdrawal of LIF induces a spontaneous heterogenous differentiation profile, enabling the cells to differentiate to all three germ layers^35^. We confirmed the AN3-12 cells’ differentiation capacity by assessing morphological changes and measuring the reduced expression of core pluripotency markers by Western blot (WB) analysis (Figure 1B, C, D; Supplementary Figure 1A). Next, we performed proteomic analysis of pluripotent cells compared to D1, D3, D5 and D7 cells to further assess successful differentiation using our protocol. Principal component analysis (PCA) revealed a cluster formation of D0 and D1 cells, a clear shift for proteins uncovered from D3 cells and another cluster consisting of D5 and D7 cells, suggesting robust changes in protein abundance during differentiation (Supplementary Figure 1B). Expression levels of different lineage markers (FGF8: primitive streak, FGF5: neuroectoderm, EOMES: endoderm, NKX2-5: mesoderm) for the same time points were measured by qPCR and confirmed the spontaneous differentiation of AN3-12 cells into all three germ layers (Figure 1E). In line with these findings, many core pluripotency markers were strongly downregulated over time revealing a loss of pluripotency (Figure 1F). Given their low abundance, the expression of these marker proteins was not detectable in our proteomics dataset.

**Figure 1:**
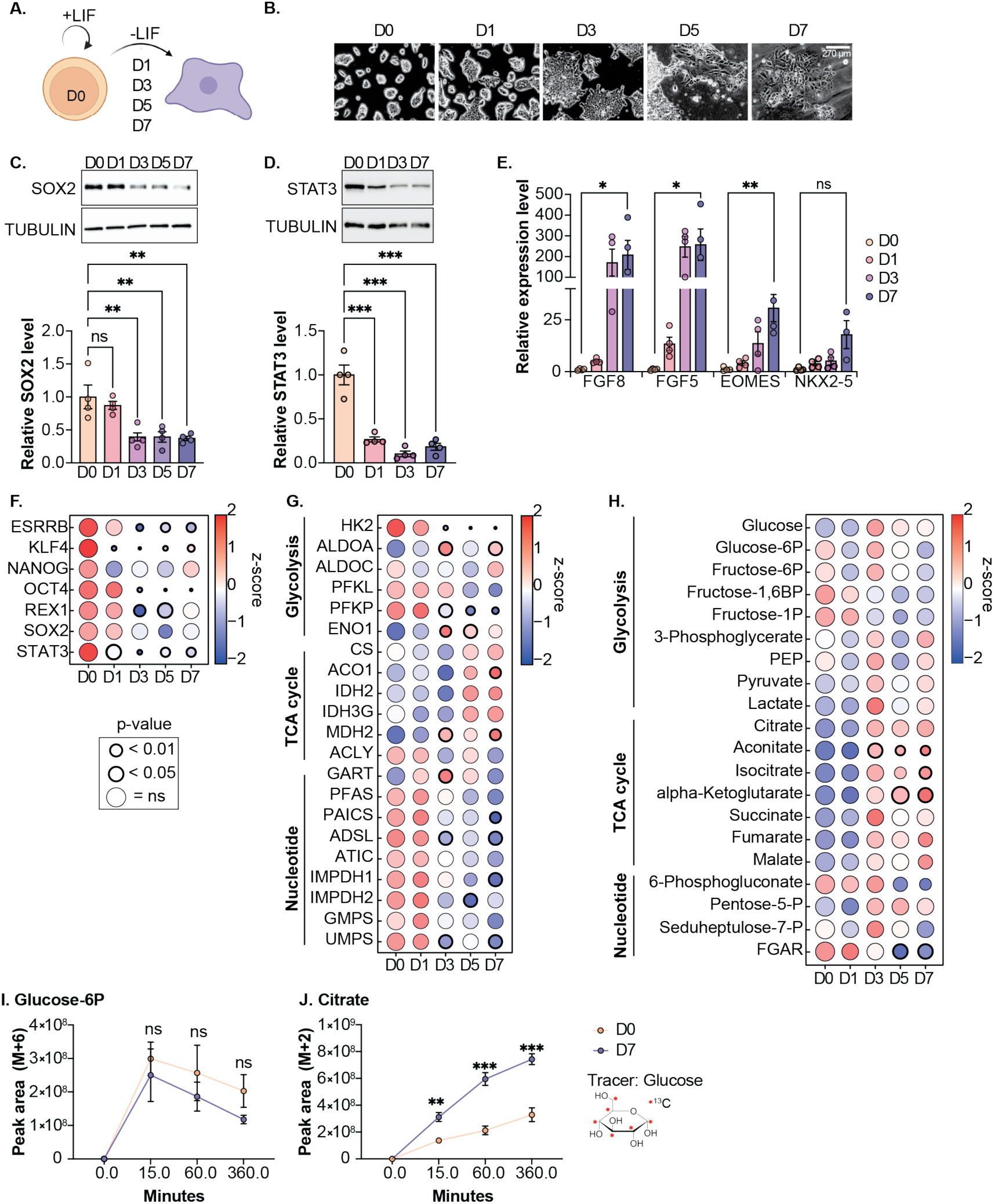
AN3-12 mouse embryonic stem cells display metabolic rewiring upon the removal of the leukaemia inhibitory factor. **A.** Schematic depiction of the experimental workflow to induce differentiation of AN3-12 mESCs by removing LIF for the specified number of days (D1–7). Pluripotent cells were cultured in medium containing LIF (D0). **B.** Representative pictures of AN3-12 mESCs during the differentiation process. Scale bar: 270 µm**. C.** Western blot analysis and quantification of SOX2 in WT cells during differentiation (mean ± SEM, n=4, ** p<0.01, One-way ANOVA Dunnett post-test). **D.** Western blot analysis and quantification of STAT3 in WT cells during differentiation (mean ± SEM, n=4, *** p<0.001, One-way ANOVA Dunnett post-test). **E.** Relative mRNA expression levels of various lineage markers in WT cells before and after differentiation (mean ± SEM, n≤4, * p<0.05, ** p<0.01, unpaired, one-tailed t-test). **F.** Proteomics data of pluripotency markers in WT cells during differentiation. Significances are indicated by differences in circle size and thickness (n=4) **G.** Proteomics data of metabolic enzymes in WT cells during differentiation. Significances are indicated by differences in circle size and thickness (n=4). **H.** Metabolomics data of different pathways in WT cells during differentiation. Significances are indicated by differences in circle size and thickness (n=4). **I.** Pluripotent and 7-day differentiated (D7) WT cells were labelled with ^13^C_6_-glucose. Shown is the labelled fraction of glucose-6-phosphate (M+6) at 0, 15, 60, and 360 minutes of labelling (mean ± SEM, n=4, ** p<0.01, *** p<0.001, Two-way ANOVA Šídák’s post-test). **J.** Same setup as in I. for citrate.

Functionally, the proteomics dataset unravelled a profound metabolic switch characterized by a downregulation of glycolytic and nucleotide metabolism-related enzymes, while enzymes regulating the TCA cycle were overall upregulated during differentiation, as described before in SCs^36^ (Figure 1G).

Metabolomics data generated from the same samples further strengthened the observation of a severe metabolic rewiring during spontaneous differentiation induced by LIF deprivation. The PCA of the metabolomics samples showed similar clustering as for the proteomics samples, with a cluster consisting of D0 and D1 cells, a separated heterogenous intermediate cluster for metabolites from D3 cells and a third cluster from D5 and D7 cells (Supplementary Figure 1C). As for the corresponding enzymes, metabolites of the TCA cycle were upregulated, while glycolytic metabolites and metabolites of the nucleotide metabolism seemed predominantly downregulated (Figure 1H). In addition to steady state measurements, we also performed metabolic isotope tracing experiments using uniformly labelled ^13^C_6_-glucose in pluripotent cells and compared them to D7 cells. Consistently, glucose uptake into glycolysis was mildly decreased during differentiation (Glc6P, Figure 1I), while ^13^C enrichment in TCA cycle intermediates were significantly upregulated (citrate, Figure 1J; alpha-ketoglutarate, Supplementary Figure 1D and succinate, Supplementary Figure 1E). Collectively, these data imply that LIF withdrawal effectively and robustly induces spontaneous differentiation of AN3-12 mESCs, initiating a substantial metabolic transition and making this model suitable for uncovering novel metabolic signatures.

### Extensive metabolic rewiring of the hexosamine biosynthetic pathway during stem cell differentiation

In a next step, we honed in on enzymes involved in the HBP (Figure 2A) because of the striking changes in expression levels detected during differentiation, as shown by qPCR (Supplementary Figure 2A) and proteomic analysis, and confirmed by WB (Figure 2B, D, E; Supplementary Figure 2B). Comparing pluripotent cells with D7 cells, we observed an enzymatic switch from GFPT2 to GFPT1 in the forward reaction with a simultaneous switch from GNPDA2 to GNPDA1 in the reverse reaction upon differentiation (Figure 2B). Furthermore, GNPNAT1 was strongly downregulated, while AMDHD2 expression remained unchanged (except for a slight decrease at D3 and D5, Supplementary Figure 2B). Milder but significant changes were observed for PGM3 and UAP1, with the former being downregulated and the latter being upregulated (Figure 2B).

**Figure 2:**
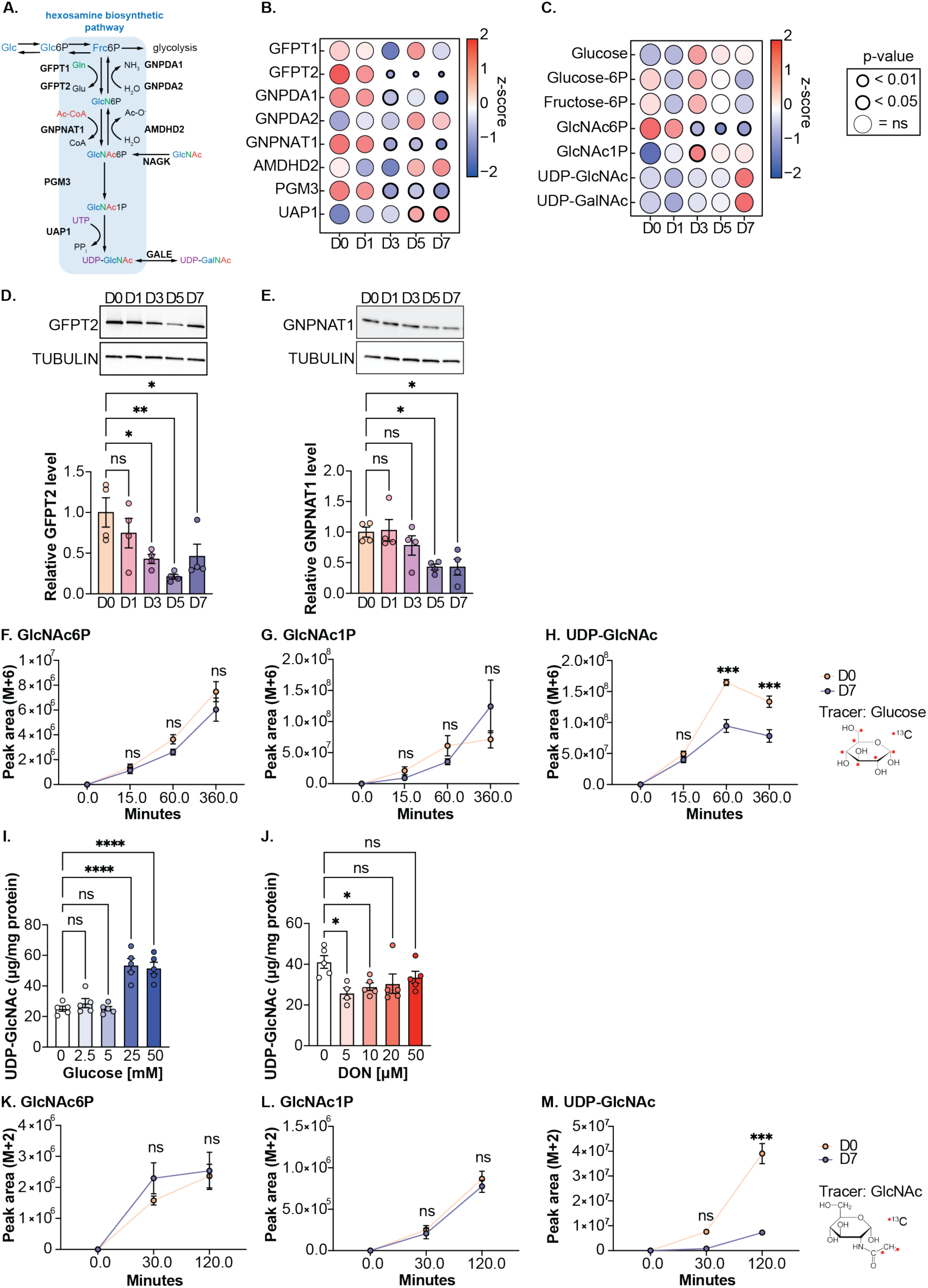
Extensive metabolic rewiring of the hexosamine biosynthetic pathway during stem cell differentiation. **A.** Schematic overview of the hexosamine pathway (blue box). The intermediate fructose-6-phosphate (Frc6P) from glycolysis is converted to UDP-GlcNAc, which is a precursor for glycosylation reactions. The enzymes are glutamine fructose-6-phosphate amidotransferase (GFPT1/2), glucosamine-6-phosphate N-acetyltransferase (GNPNAT1), phosphoglucomutase (PGM3), UDP-N-acetylglucosamine pyrophosphorylase (UAP1), glucosamine-6-phosphate deaminase (GNPDA1/2), N-acetylglucosamine deacetylase (AMDHD2), and UDP-galactose-4’-epimerase (GALE). **B.** Proteomics data of the HBP enzymes in WT cells during differentiation. Significances are indicated by differences in circle size and thickness (n=4). **C.** Metabolomics data of the HBP intermediates in WT cells during differentiation. Significances are indicated by differences in circle size and thickness (n=4). Glucose, glucose-6P and fructose-6P data are taken from Figure 1H. **D.** Western blot analysis and quantification of GFPT2 in WT cells during differentiation (mean ± SEM, n=4, * p<0.05, ** p<0.01, One-way ANOVA Dunnett post-test). **E**. Western blot analysis and quantification of GNPNAT1 in WT cells during differentiation (mean ± SEM, n=4, * p<0.05, One-way ANOVA Dunnett post-test). **F.** Pluripotent and D7 WT cells were labelled with ^13^C_6_-glucose. Shown is the labelled fraction of GlcNAc6P (M+6) at 0, 15, 60, and 360 minutes of labelling (mean ± SEM, n=4, *** p<0.001, Two-way ANOVA Šídák’s post-test). **G.** Same setup as in F. for GlcNac1P and **H.** UDP-GlcNAc. **I.** IC-MS analysis of UDP-GlcNAc levels in WT cells cultured for 24 h in medium containing 0, 2.5, 5, 25, or 50 mM glucose (mean ± SEM, n=5, **** p<0.0001, One-way ANOVA Tukey post-test). **J.** IC-MS analysis of UDP-GlcNAc levels in WT cells cultured for 24 h in medium containing different concentration of the GFAT inhibitor DON (mean ± SEM, n=5, * p<0.05, One-way ANOVA Tukey post-test). **K.** Pluripotent and D7 WT cells were labelled with 10 mM ^13^C_2_-GlcNAc. Shown is the labelled fraction for GlcNAc6P(M+2) at 0, 30, and 120 minutes of labelling (mean ± SEM, n=4, *** p<0.001, Two-way ANOVA Šídák’s post-test). **L.** Same setup as in K. for GlcNac1P and **M.** for UDP-GlcNAc.

To investigate the consequences of this metabolic rewiring during stem cell differentiation, we analyzed the HBP intermediates in the metabolomics dataset of the same samples. Surprisingly, we observed only significantly decreased GlcNAc6P steady-state levels in D7 cells compared to D0 cells, while GlcNAc1P and UDP-GlcNAc/UDP-GalNAc levels were slightly but not significantly increased (Figure 2C). Upstream metabolites including intracellular glucose, Glc6P and Frc6P levels were not significantly changed (Figure 2C). When examining the glucose enrichment through the HBP using uniformly-labelled ^13^C_6_-glucose, we observed no drastic differences for labelled GlcNAc6P (Figure 2F) and GlcNAc1P (Figure 2G) when comparing pluripotent and D7 cells, while significantly less labelled UDP-GlcNAc was generated (Figure 2H).

Next, we assessed O-GlcNAcylation to test whether the reduced synthesis of UDP-GlcNAc results in lower levels of this relevant post-translational modification (PTM). Despite the reduced ^13^C_6_ glucose-mediated enrichment into UDP-GlcNAc (Figure 2H), no significant changes in global O-GlcNAcylation levels were observed in the WB analysis (Supplementary Figure 2C), which matches the steady-state UDP-GlcNAc levels (Figure 2C). Interestingly, OGT expression, which regulates O-GlcNAcylation and in turn stabilization of the pluripotency factors SOX2 and OCT4, is reduced during differentiation, while OGA levels remain stable (Supplementary Figure 2D), suggesting that specific targets may be regulated differently to the overall PTM levels^23^. Given the decreased expression of relevant HBP enzymes and the reduced glucose enrichment towards UDP-GlcNAc, but without a corresponding decrease in UDP-GlcNAc steady-state levels and total O-GlcNAcylation, we sought to determine whether glucose is the sole source for the production of UDP-GlcNAc in pluripotent cells. To diminish *de novo* synthesis, we cultivated cells in media containing different glucose concentrations before measuring UDP-GlcNAc. Indeed, higher glucose concentrations (25 mM and 50 mM glucose) resulted in significantly increased UDP-GlcNAc compared to lower substrate levels (5 mM glucose) (Figure 2I). Surprisingly, we observed similar UDP-GlcNAc levels at 2.5 mM or 5 mM glucose as in WT cells cultured for 24 hours in glucose-free medium (Figure 2I), suggesting the presence of an alternative source for UDP-GlcNAc production in the absence of glucose. In addition to limiting the substrate of the HBP, we tested varying concentrations of the GFPT1/2 inhibitor 6-diazo-5-oxo-norleucine (DON) to block *de novo* synthesis prior to measuring UDP-GlcNAc levels. Despite inhibition of the rate-limiting enzymes, we observed only a reduction of less than 50%, further suggesting the presence of an alternative synthesis pathway (Figure 2J). It has been described in cancer cells that under conditions of limited nutrient availability, free GlcNAc released from degraded proteins can be phosphorylated via the kinase NagK to enter the HBP as GlcNAc6P, thereby supplying the UDP-GlcNAc pool independently of glucose^37^. This process is referred to as the salvage pathway. However, whether this also applies to pluripotent mESCs and if this salvage pathway is regulated differently in differentiating cells has not yet been investigated. To test this hypothesis, we supplemented pluripotent and differentiated cells (D7) with ^13^C_2_(acetate)-labelled GlcNAc to trace its enrichment into the HBP. Surprisingly, GlcNAc entered the HBP in pluripotent and in D7 cells despite unlimited glucose availability, as can be seen by similar levels of ^13^C_2_-labelled GlcNAc6P (Figure 2K) and GlcNAc1P (Figure 2L). Similar as for the *de novo* synthesis pathway, the continued flux into UDP-GlcNAc was significantly decreased in differentiated cells (Figure 2M).

Taken together, these data suggest a profound metabolic rewiring of the HBP during SC differentiation, with a downregulation of many of the enzymes catalyzing the forward reactions. We further showed that pluripotent mESCs rely on a combination of the *de novo* synthesis and the GlcNAc salvage pathway to maintain UDP-GlcNAc levels.

### *De novo* and salvage pathway-supplied GlcNAc6P is readily converted to Glucosamine-6-phosphate by AMDHD2 in pluripotent and differentiated cells

Since pluripotent SCs rely on both *de novo* synthesis and GlcNAc salvage to maintain their UDP-GlcNAc pool, we sought to better understand the role of GlcNAc6P, the metabolite where these pathways converge. Its central position, along with its marked downregulation during differentiation, prompted us to consider GlcNAc6P as a key candidate for further investigation. To test if manipulation of GlcNAc6P levels influences SC fate, we first knocked down enzymes relevant for GlcNAc6P metabolism of the *de novo* synthesis (GNPNAT1 and AMDHD2) in pluripotent ESCs. The siRNA-mediated knockdown (KD) of GNPNAT1 and AMDHD2 was efficient as confirmed by WB analysis, resulting in 81% and 78% reduction in protein levels, respectively, when compared to control cells treated with a non-targeting siRNA (Supplementary Figure 3A). While none of the measured HBP intermediates showed a significant change in their steady-state levels after knockdown of GNPNAT1 or AMDHD2 (Supplementary Figure 3B, C, D), stable isotope tracing with ^13^C_6_-glucose revealed a significant and 4-fold higher labelling of GlcNAc6P in the AMDHD2 siRNA-treated cells only when compared to control cells (Figure 3B). This suggests an accumulation of this metabolite due to a disturbed reverse flux, assuming similar glucose uptake as evidenced by similar degrees of glucose-6P labelling (Figure 3A). The response to AMDHD2 KD appears to be very local and specific seeing as none of the conditions tested showed a significant change in the levels of any of the other labelled HBP intermediates (GlcNAc1P, Figure 3C and UDP-GlcNAc, Figure 3D) or in those of related pathways such as glycolysis (Glc6P, Figure 3A), PPP (Supplementary Figure 3E), the TCA cycle (Supplementary Figure 3F) or glutamine metabolism (Supplementary Figure 3G). Since the deacetylation of GlcNAc6P to GlcN6P catalyzed by AMDHD2 significantly limits GlcNAc6P levels, we used AMDHD2 knock-out (KO) AN3-12 cells for further analyses^32^. We compared the metabolic profiles of the AMDHD2 KO cells with control cells and found significantly higher steady-state levels for all measured HBP intermediates (Frc6P, Figure 3F; GlcNAc6P, Figure 3G; GlcNAc1P, Figure 3H and UDP-GlcNAc, Figure 3I), further confirming the previously observed increase in UDP-GlcNAc levels^32^. The most drastic change was observed for GlcNAc6P with a 104-fold enrichment in the AMDHD2 KO line compared to the WT, emphasizing the importance of AMDHD2 for GlcNAc6P catabolism. Interestingly, when examining the expression levels of other HBP enzymes in KO versus WT cells, we observed a mild, though not significant, downregulation of the enzymes responsible for the forward reaction (GFPT1/2, UAP1), alongside an upregulation of the counteracting enzymes (GNPDA1/2), which may help compensate for the increased UDP-GlcNAc levels (Supplementary Figure 3H). While no significant differences were detected in steady-state metabolite levels in other related pathways (Supplementary Figure 3J), the AMDHD2 KO cells show downregulation of some enzymes of the nucleotide metabolism and relative overexpression of most enzymes of glycolysis (HK2, ALDOA, PFKL, ENO1, Supplementary Figure 3I), suggesting a compensatory response to the absent back-flux into Fr6P.

**Figure 3:**
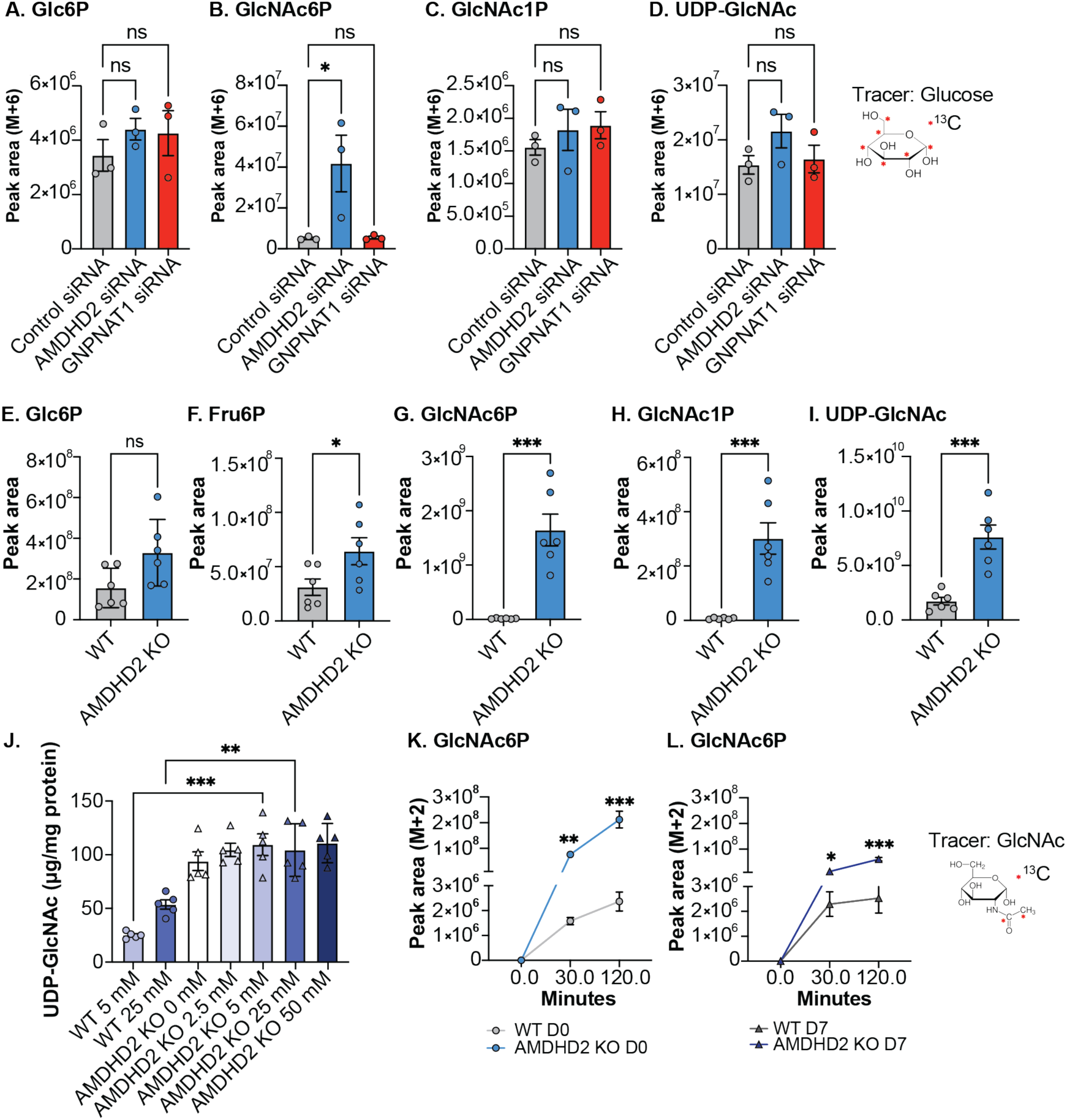
*De novo* and salvage pathway-supplied GlcNAc6P is readily converted to Glucosamine-6-phosphate by AMDHD2 in pluripotent and differentiated cells. **A.** mESCs were treated with non-targeting siRNAs or siRNAs targeting AMDHD2 or GNPNAT1, respectively. After 48 h, cells were labelled with ^13^C_6_-glucose. Shown is the labelled fraction of glucose-6-P (M+6) at 15 minutes of labelling (mean ± SEM, n=3, * p<0.05, One-way ANOVA Dunnett post-test). **B.** Same setup as in A. for GlcNAc6P, **C.** for GlcNAc1P and **D.** UDP-GlcNAc. **E.** IC-MS analysis of Glc6P in WT and AMDHD2 KO cells (mean ± SEM, n=6, * p<0.05, *** p<0.001, unpaired, one-tailed t-test). **F.** Same setup as in E. for Fru6P, **G.** for GlcNAc6P, **H.** for GlcNAc1P and **I.** for UDP-GlcNAc. **J.** IC-MS analysis of UDP-GlcNAc levels in WT and AMDHD2 KO cells cultured for 24 h in medium containing 0, 2.5, 5, 25, or 50 mM glucose (mean ± SEM, n=5, ** p<0.01, *** p<0.001, One-way ANOVA Tukey post-test). **K.** Pluripotent WT and AMDHD2 KO mESCs were labelled with 10 mM ^13^C_2_-GlcNAc. Shown is the labelled fraction of GlcNAc6P (M+2) at 0, 30, and 120 minutes of labelling (mean ± SEM, n=4, ** p<0.01, *** p<0.001, Two-way ANOVA Šídák’s post-test). **L.** Same setup as in K for D7-differentiated WT and AMDHD2 KO mESCs.

Surprisingly, when we compared the contributing factor of the *de novo* synthesis and the salvage pathway in producing UDP-GlcNAc in AMDHD2 KO cells, we found that when AMDHD2 KO cells were cultured in glucose-free medium, UDP-GlcNAc levels were stably enriched as in medium with high glucose concentrations (Figure 3J). This suggests a reduced dependency on the *de novo* synthesis for UDP-GlcNAc production in the absence of AMDHD2. In accordance, when using labelled GlcNAc to trace the salvage pathway in pluripotent WT and AMDHD2 KO cells, we detected dramatically increased levels of ^13^C_2_-labelled GlcNAc6P in KO cells (Figure 3K). While the labelled fraction of GlcNAc6P did not change after seven days of differentiation in WT cells, the levels in the KO were decreased in D7 compared to pluripotent KO cells but remained significantly higher than in WT cells following differentiation (Figure 3L). These data suggest a reduced AMDHD2 or NagK activity at D7. Interestingly, despite the significant accumulation of GlcNAc6P in the AMDHD2 KO in both pluripotent and D7 cells, the levels of labelled UDP-GlcNAc were markedly reduced compared to the corresponding WT control cells, suggesting potential feedback regulation mediated by UAP1 (at D0) and PGM3 (at D7) (Supplementary Figure 3H, M, N). Notably, when we measured steady-state levels in pluripotent KO cells compared to D7 cells, we observed a similar metabolic shift as in WT cells, accompanied by a slight decrease in glycolytic metabolites (Supplementary Figure 3O–Q), whereas TCA cycle intermediates were markedly increased after differentiation (Supplementary Figure 3R, S). However, the HBP intermediates where unchanged and persisted at high levels after seven days of differentiation (GlcNAc6P, Supplementary Figure 3T; GlcNAc1P, Supplementary Figure 3U; UDP-GlcNAc, Supplementary Figure 3V). Regardless of the dramatic accumulation of total UDP-GlcNAc levels in the KO, downstream O-GlcNAcylated protein levels remained stable (Supplementary 3W), probably due to the significant downregulation of the corresponding enzyme OGT (Supplementary Figure 3Y). Collectively, these findings suggest that AMDHD2 is a critical factor in maintaining the continuous deacetylation of GlcNAc6P, supplied through *de novo* synthesis and the salvage pathway, and that it plays a particularly important role in pluripotent cells.

### The loss of AMDHD2 in mESCs impairs differentiation into all germ layers

Next, we investigated the effect of AMDHD2 KO on pluripotency exit and SC differentiation since AMDHD2 appears to be particularly important in pluripotent cells. Spontaneous differentiation through LIF withdrawal for seven days resulted in substantial increases in lineage marker expression in WT cells, whereas expression of the same genes in the AMDHD2 KO cells was markedly lower, indicating a compromised differentiation (BRACHYURY: primitive streak; FGF5: neuroectoderm; HAND1: mesoderm; EOMES: endoderm; Figure 4A–D). Notably, the mRNA expression levels of pluripotency factors were not elevated in the KO compared to WT cells, either in the pluripotent state or throughout differentiation (TCL1, Supplementary Figure 4A; KLF4, Supplementary Figure 4B; SOX2, Supplementary Figure 4C). In fact, the protein levels of the pluripotency factors appear rather downregulated in pluripotent KO compared to WT cells (Supplementary Figure 4D). To avoid potential genetic compensation mechanisms that might mask primary effects caused by the severe intervention of a complete KO, we generated a mESC line allowing conditional depletion of AMDHD2 using the Auxin-Inducible Degron (AID) system integrated by CRISPR/Cas9 gene editing (Figure 4E)^38^. Efficient depletion of AMDHD2 occurred within 1 hour (h) after auxin/ indole-3-acetic acid (IAA) treatment and in a time- and concentration-dependent manner (Supplementary Figure 4E). Treatment with 0.5 mM IAA for 6 h was sufficient to decrease AMDHD2 levels to 8 % (Supplementary Figure 4E, G), while IAA treatment in control WT cells only harbouring the OsTIR1-V5 ligase but no AID-tagged target protein had no effect on the protein abundance of AMDHD2 (Supplementary Figure 4F). When IAA was removed after 6 h of depletion, AMDHD2 levels returned to untreated levels after 72 h (Figure 4F). After depleting the target protein in AMDHD2-AID mESCs for 6 h, the cells were subjected to differentiation by LIF deprivation as previously described without further addition of IAA. Consistent with AMDHD2 KO, we observed a similarly compromised differentiation in AMDHD2-depleted cells compared to control cells, impacting all germ layers (Figure 4G–J). To evaluate the functional loss, we measured HBP intermediates in AMDHD2-AID cells at various time points following 6 h of depletion, observing the most pronounced effect after 48 h of recovery. While GlcNAc6P (Supplementary Figure 4H) and GlcNAc1P (Supplementary Figure 4I) were significantly elevated after IAA treatment, UDP-GlcNAc levels remained unchanged (Supplementary Figure 4J). Thus, the data highlight the critical role of AMDHD2 during SC differentiation, as AMDHD2’s loss significantly impaired the differentiation potential.

**Figure 4:**
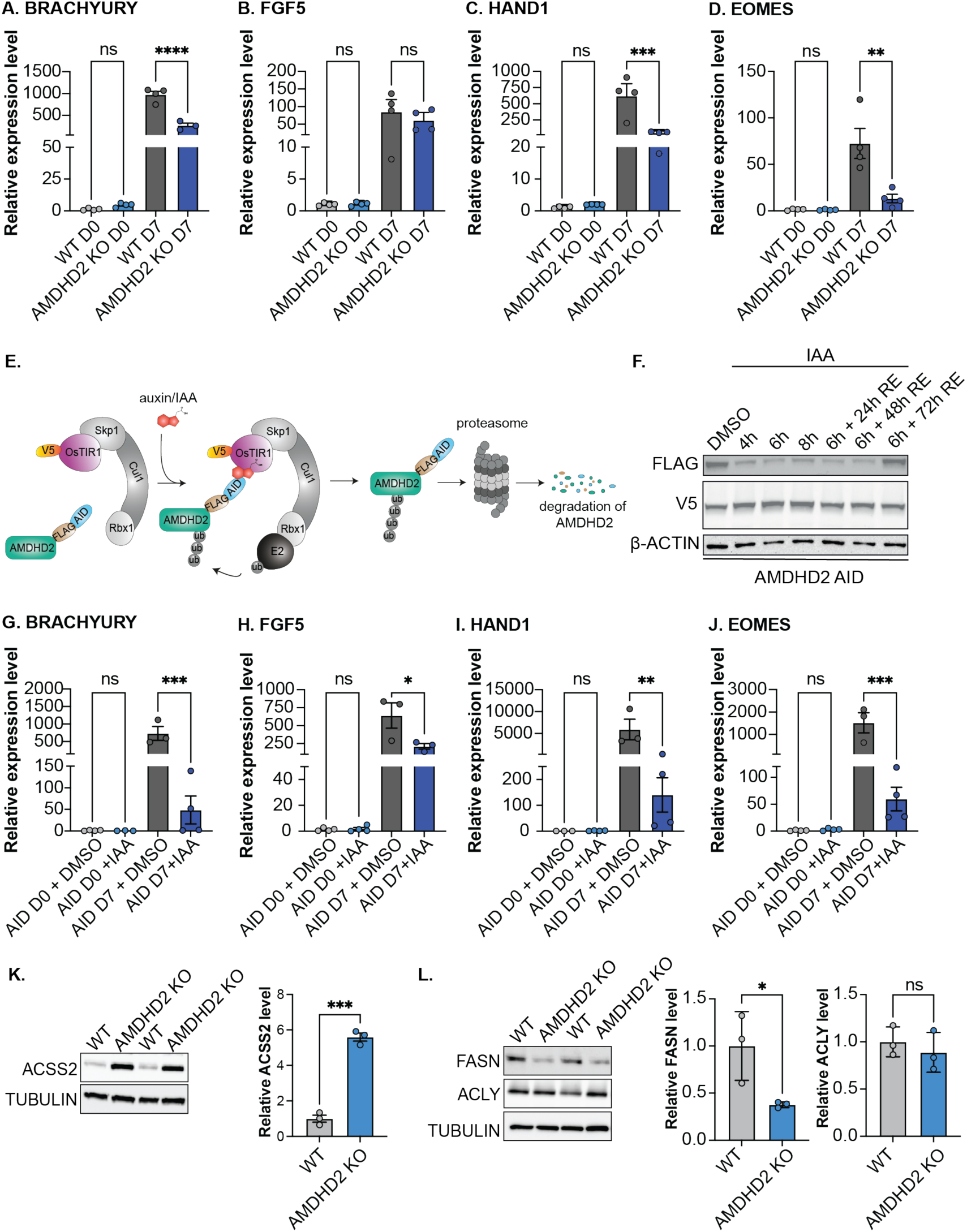
The loss of AMDHD2 in mESCs impairs differentiation into all germ layers. **A.** Relative mRNA expression of BRACHYURY in pluripotent and 7-day differentiated WT and AMDHD2 KO cells (mean ± SEM, n=4, ** p<0.01, *** p<0.001, **** p<0.0001, One-way ANOVA Tukey post-test). **B.** Same setup as in A. for FGF5, **C.** for HAND1, **D.** for EOMES. **E.** Schematic representation of the Auxin-inducible degron (AID) system. The addition of auxin/IAA promotes the binding of the AID motif to Transport Inhibitor Response 1 (TIR1), which recruits the Skp1-Cullin-F-box (SCF) complex, leading to ubiquitination and subsequent proteasomal degradation. To this end, we introduced the Oryza sativa TIR1 (OsTIR1) E3 ligase fused to an V5-tag into the TIGRE locus of WT mESCs and additionally tagged endogenous AMDHD2 with a C-terminal FLAG-AID-tag. **F.** Western Blot analysis to confirm functionality of the AMDHD2-AID system. AMDHD2-FLAG-AID cells additionally expressing the OsTIR1-V5 ligase were treated with DMSO or IAA for 4, 6 or 8 h. After 6 h of IAA treatment, IAA was removed and cells were allowed to recover for 24, 48 and 72 h. **G.** Relative mRNA expression of BRACHYURY in pluripotent and D7 AMDHD2-AID cells. The cells were treated with DMSO or 0.5 mM IAA for 6 h before induction of differentiation (mean ± SEM, n=3, * p<0.05, ** p<0.01, *** p<0.001, One-way ANOVA Tukey post-test). **H.** Same setup as in G. for FGF5, **I.** for HAND1, **J.** for EOMES. **K.** Western blot analysis and quantification of ACSS2 in pluripotent WT and AMDHD2 KO cells (mean ± SD, n=3, * p<0.05, *** p<0.001, unpaired, one-tailed t-test). **L.** Same setup as in K. for FASN and ACLY.

Based on the compromised differentiation of SCs upon AMDHD2 loss, we aimed to explore the specific mechanism through which AMDHD2 activity controls differentiation. Notably, IAA-induced AMDHD2 depletion did not increase UDP-GlcNAc levels (Supplementary Figure 4J), implying a pathway that operates independently of the HBP end product. Relevantly, AMDHD2 catalyzes the deacetylation of GlcNAc6P, which led us to hypothesize that the released acetate could be converted to acetyl-CoA through acetyl-CoA synthetase 2 (ACSS2), thereby regulating the nucleo-cytosolic acetyl-CoA pool. It is well-established that intracellular acetate and acetyl-CoA levels are linked to SC maintenance, with their availability playing a crucial role in regulating lipid metabolism, chromatin accessibility, histone acetylation, and, consequently, gene expression profiles^7,39^. Previous studies have highlighted the importance of maintaining stable acetyl-CoA levels by demonstrating a compensatory upregulation of ACSS2 in various cancer cell lines and MEFs when ACLY is disrupted^8–11^. To this end, we checked the expression of relevant metabolic enzymes in our proteomics data of our AMDHD2 KO cells (Supplementary Figure 4K–M) and by WB analysis (Figure 4K, L). We found that ACSS2 is significantly upregulated in these cells (Figure 4K, Supplementary Figure 4K), while FASN is significantly downregulated (Figure 4L, Supplementary Figure 4L). Of note, ACLY levels were unchanged (Figure 5L, Supplementary Figure 4L). Together, these data suggest that loss of AMDHD2 might severely interfere with acetate and the connected acetyl-CoA metabolism.

**Figure 5:**
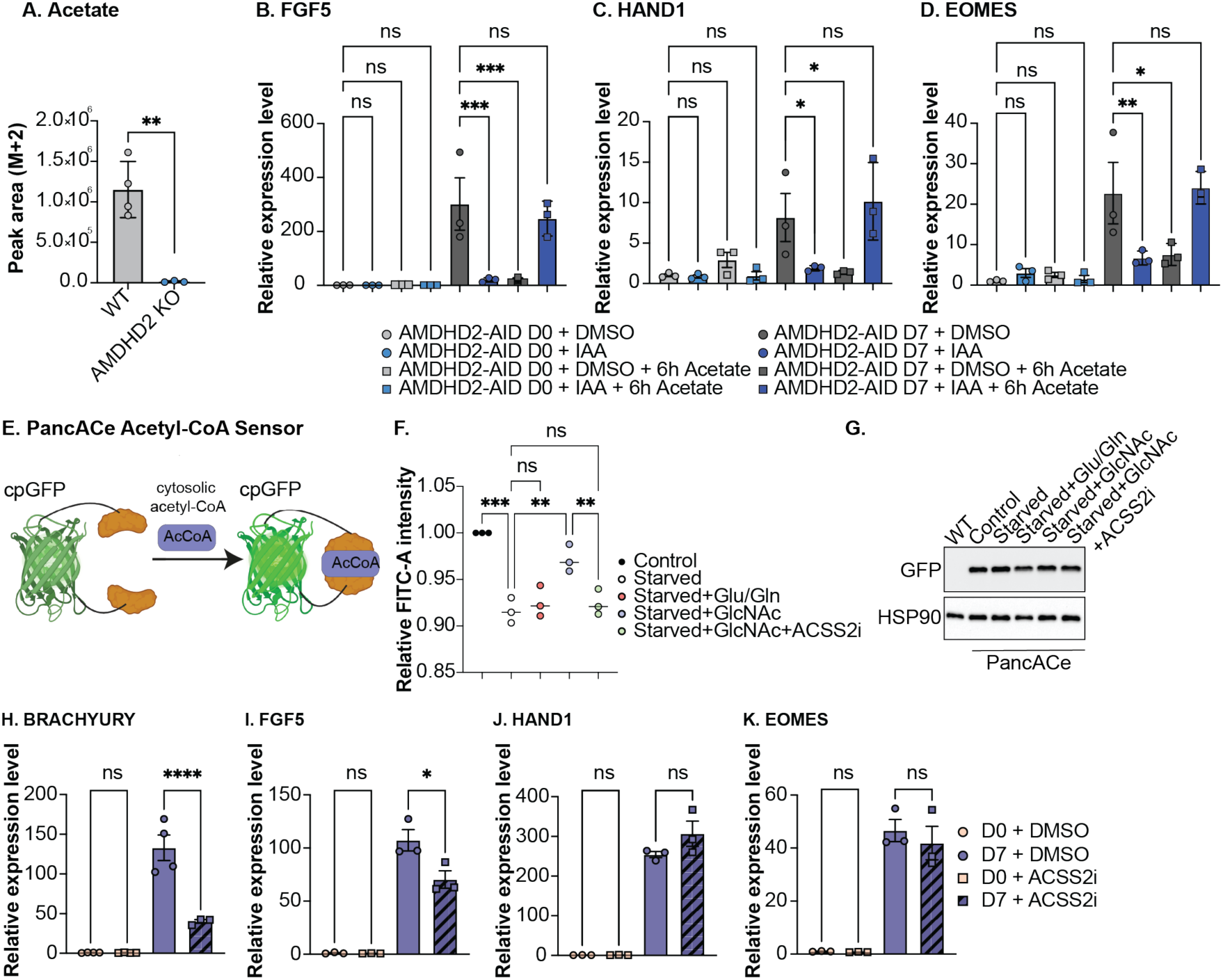
AMDHD2 impacts stem cell differentiation by altering acetate levels in the cell. **A.** Measurement of labelled acetate (M+2) in WT or AMDHD2 KO cells, after treatment with 10 mM ^13^C_2_-GlcNAc for 48 h and simultaneous incubation with 10 µM ACSS2 inhibitor (mean ± SEM, n≥4, unpaired, one-tailed t-test). **B.** Relative mRNA expression of FGF5 in pluripotent and D7 WT cells. Cells were treated with either DMSO or IAA for 6 hours with or without 5 mM acetate (mean ± SEM, n=3, * p<0.05, ** p<0.01, *** p<0.001, Two-way ANOVA Tukey post-test). **C.** Same setup as in B. for HAND1, in **D.** for EOMES. **E.** Schematic depiction of the fluorescent biosensor for cytosolic acetyl-CoA (AcCoA). Binding of AcCoA induces a conformational change in the circularly permuted GFP (cpGFP) sensor, triggering fluorescence^40^. **F.** Relative fluorescent signal of WT cells stably expressing the cytosolic acetyl-CoA sensor. Control cells were cultivated in regular stem cell medium while starved cells were incubated for 4 h in medium lacking the precursors glucose and glutamine. For 1 h, cells were either re-fed with regular medium containing glucose and glutamine, with medium containing only GlcNAc or GlcNAc and 10 μM ACSS2 inhibitor (mean ± SD, n=3, ** p<0.01, *** p<0.001, One-way ANOVA Tukey post-test). **G.** Western blot analysis of AcCoA-expressing mESCs from F. using a GFP antibody. **H.** Relative mRNA expression of BRACHYURY in pluripotent and D7 WT cells. Cells were treated with DMSO or 1 µM ACSS2 inhibitor for 24 hours at D0 or with DMSO or 0.1 µM ACSS2 inhibitor for the first 72 h of differentiation (mean ± SEM, n=3, * p<0.05, **** p<0.0001, One-way ANOVA Tukey post-test). **I.** Same setup as in H. for FGF5, in **J.** for HAND1 and in **K.** for EOMES.

### AMDHD2 impacts stem cell pluripotency by altering cytosolic acetate levels in the cell

Next, we aimed to determine whether AMDHD2 impacts acetate levels in cells, as suggested by the adaptations observed in AMDHD2 KO cells (Figures 4K, L). To this end, we utilized ^13^C_2_(acetate)-labelled GlcNAc in WT and AMDHD2 KO cells and measured labelled acetate levels. A labelling time of 48 hours, together with ACSS2 inhibition, was necessary to enable a robust measurement of labelled acetate. Compared to WT cells, we could only detect trace amounts of labelled acetate in AMDHD2 KO cells (Figure 5A), which supports the assumption that AMDHD2 utilizes GlcNAc6P to generate acetate. Although there was no significant difference in steady-state acetate levels between AMDHD2 KO and WT cells (Supplementary Figure 5A), acutely depleting AMDHD2 in AMDHD2-AID cells for 6 h resulted in a fivefold reduction in steady-state acetate levels (Supplementary Figure 5B). We hypothesized that if reduced acetate levels were critical to the compromised differentiation of AMDHD2 KO and AID cells, then supplementing acetate during the 6h AMDHD2-depletion period would rescue differentiation in the AID cells. Indeed, differentiation was restored to the WT differentiation pattern (Figure 5B–D). However, acetate supplementation in cells where AMDHD2 was not depleted also resulted in compromised differentiation, as reported previously^7^. This indicates that both insufficient and excessive acetate levels negatively impact differentiation. Surprisingly, acetate supplementation resulted in a more effective rescue than allowing AMDHD2-depleted cells to recuperate for 72 h, during which time AMDHD2 protein levels have recovered (Figure 4F), before differentiation (Supplementary Figure 5C–E). The metabolic adaptations undergone by the AID cells through AMDHD2 depletion appear to require more time to reset, suggesting that potential downstream effects may be responsible for the differentiation phenotype.

Therefore, we investigated if GlcNAc6P deacetylation can affect subcellular acetyl-CoA levels. We used a previously developed acetyl-CoA sensor specifically designed to detect cytosolic acetyl-CoA levels (Figure 5E)^40^. After introducing this sensor into mESCs, we induced a reduction in cytosolic acetyl-CoA levels by depleting the precursor molecules glucose and glutamine for 4 h (Figure 5F). Adding both substrates back for 1 h was insufficient to recover acetyl-CoA levels. However, refeeding the cells with a medium containing GlcNAc as the sole carbon source for 1 h was sufficient to restore cytosolic acetyl-CoA levels. When ACSS2 was inhibited alongside the supply of GlcNAc, acetyl-CoA levels remained reduced. We verified the expression of the stably expressed acetyl-CoA sensor in the WT AN3-12 cells using Western blotting with a GFP antibody (Figure 5G). These data confirmed that GlcNAc can be used as a source of cytosolic acetyl-CoA via ACSS2-mediated conversion of acetate to acetyl-CoA.

We further hypothesized that if a loss in acetate-supplied acetyl-CoA pools as caused by AMDHD2 KO or IAA-induced AMDHD2 depletion is a leading cause in the compromised SC differentiation, ACSS2 inhibition should mimic the compromised differentiation phenotype. In our AMDHD2-AID model, the AMDHD2 protein was only depleted for the first 72 h of differentiation (Figure 4F). Therefore, we mimicked this condition by inhibiting ACSS2 activity during the first 72 hours of differentiation in WT cells. This was sufficient to reduce the expression of lineage markers such as BRACHYURY (primitive streak; Figure 5H), FGF5 (neuroectoderm; Figure 5I), FGF8 (primitive streak; Supplementary Figure 5F), OTX2 (neuroectoderm; Supplementary Figure 5G) and MEIS1 (mesoderm; Supplementary Figure 5I) at D7. However, it did not affect the expression of HAND1 (mesoderm; Figure 5J), SOX17 (endoderm; Supplementary Figure 5H) or EOMES (endoderm; Figure 5K). While ACSS2 inhibition generally compromised differentiation somewhat similarly to AMDHD2 KO (Figure 4A–D) and AMDHD2 AID cells (Figure 4G–J), reduced ACSS2 activity may affect certain lineages more than others, such as the neuroectoderm and primitive streak. These data show that AMDHD2 catalyzes acetate release and strongly suggests that the produced acetate can be used by ACSS2 to fuel the cytosolic acetyl-CoA pool, which impacts productive stem cell differentiation.

### Loss of AMDHD2 limits lipid synthesis and modulates Histone H3K27 acetylation and methylation patterns in the promoter regions of pluripotency and differentiation genes

Cytosolic acetyl-CoA is essential for the biosynthesis of various anabolic metabolites, but measuring it is technically challenging. Therefore, we tested whether we could detect any enrichment of ^13^C_2_ in acetyl-carnitine after 24 h labelling time, as the acyl group of acetyl-CoA is readily transferred to carnitine^41^. We observed a ^13^C_2_-labelled fraction in acetyl-carnitine in WT cells, which was completely absent in untreated control cells and ^13^C_2_-GlcNAc-treated AMDHD2 KO cells (Figure 6A). To test whether the ^13^C_2_-labelled acetate could support more complex lipid synthesis downstream, we performed lipid measurements and detected AMDHD2-dependent ^13^C_2_-labelled phosphatidylcholine (Figure 6B), phosphatidylethanolamine (Figure 6C), ceramide (Figure 6D) and sphingomyelin (Figure 6E). Additionally, we observed that the steady-state levels of sphingomyelin (Supplementary Figure 6E) and cholesteryl ester (Supplementary Figure 6G) were altered, suggesting an alteration in the lipid composition of the AMDHD2 KO cells (Supplementary Figure 6B–G). This finding aligns with existing literature demonstrating that lipid composition can influence stem cell identity and pluripotency^42, 43^. Of note, we were not able to detect any enrichment of labelling in the TCA cycle metabolites after 24 h. Taken together, these data suggest that AMDHD2 activity is required during early differentiation to provide cytosolic acetate/acetyl-CoA levels by deacetylation of GlcNAc6P and in turn to maintain mESCs’ lipid composition.

**Figure 6:**
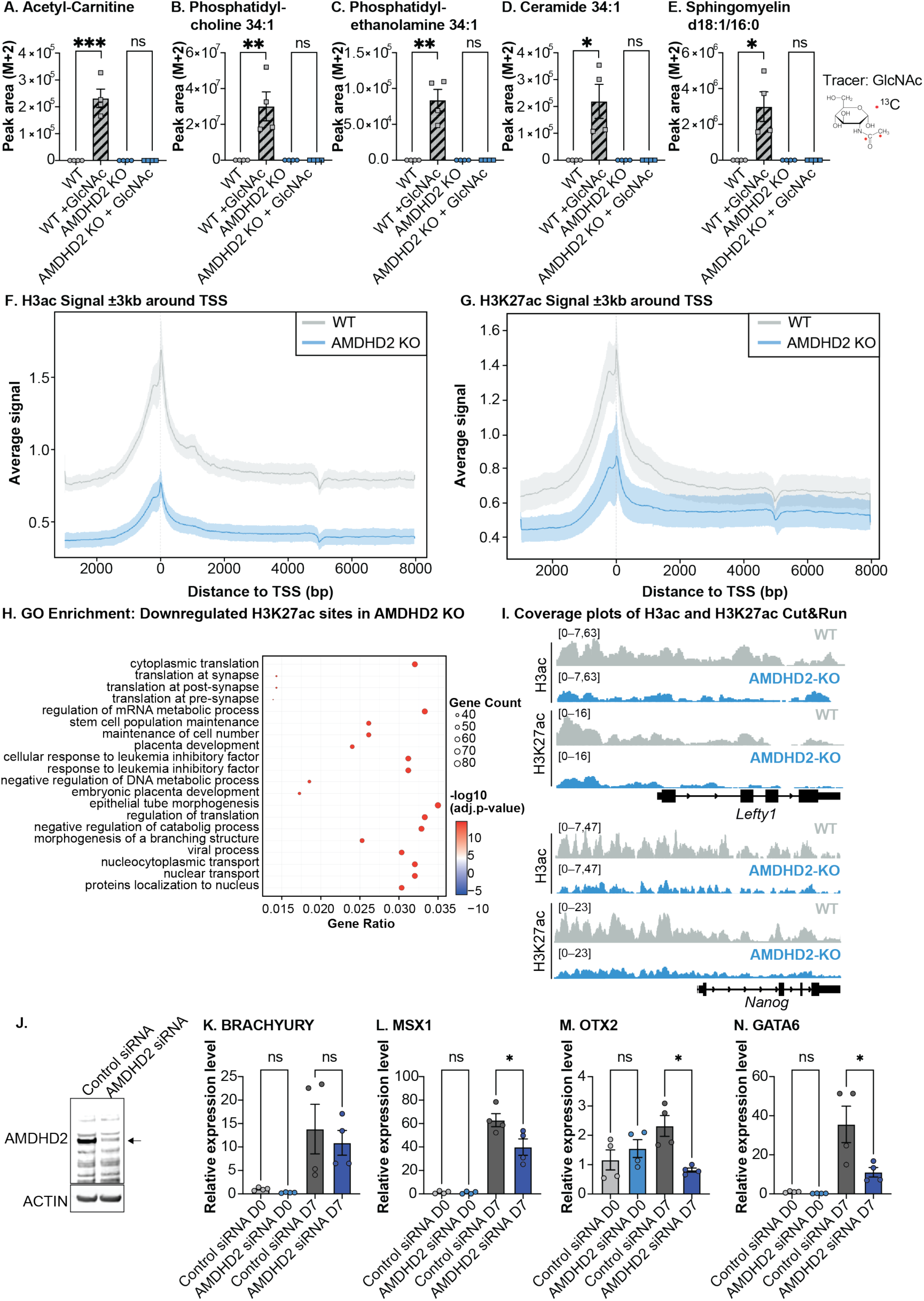
AMDHD2-mediated acetate modulates histone H3 acetylation pattern in gene promoter regions and regulates expression of pluripotency and differentiation genes in mESCs. **A.** Pluripotent WT and AMDHD2 KO mESCs were labelled with 10 mM ^13^C_2_-GlcNAc. Shown is the labelled fraction (M+2) of acetyl-carnitine after 24 h of labelling (mean ± SEM, n=4, *p<0.05, ** p<0.01, *** p<0.001, unpaired, one-tailed t-test). **B.** Same setup as in A. for phosphatidylcholine, in **C.** for phosphatidylethanolamine, in **D.** for ceramide and in **E.** for sphingomyelin. **F.** Mapping of H3acetylation (H3ac) CUT&RUN signal 3 kb upstream and downstream of the transcription start site from WT and AMDHD2 KO cells. **G.** Same setup as in F. for H3K27 acetylation signal (H3K27ac). **H.** GO-term enrichment for the genes with reduced H3K27ac signal in AMDHD2 KO cells. **I.** Coverage plots of H3ac and H3K27ac CUT&RUN data for WT and AMDHD2 KO cells surrounding the genes *Lefty1* and *Nanog*. **J.** Western blot analysis of AMDHD2 in human iPSCs 48 h post transfection with non-targeting siRNA or siRNA against AMDHD2. The expected AMDHD2 band is indicated by an arrow **K.** Relative mRNA expression of BRACHYURY in pluripotent human iPSCs and 7-day differentiated human iPSCs following treatment with control siRNA or siRNA AMDHD2 (mean ± SEM, n=4, * p<0.05, One-way ANOVA Tukey post-test). **L.** Same setup as in J. for MSX1, in **M.** for OTX2 and in **N.** for GATA6.

In view of the impact of AMDHD2 loss on the differentiation potential of mESCs and, more specifically, on acetate levels, we hypothesized that such metabolic alterations would translate into changes in the locus-specific histone acetylation landscape, thereby modulating gene expression programmes critical for SC identity and fate commitment. To test this, we performed chromatin profiling using CUT&RUN for both pan-histone H3 acetylation (H3ac) and the active enhancer/promoter mark H3K27ac (Figure 6F, G). This analysis revealed marked alterations in acetylation profiles, with distinct and reproducible binding patterns emerging in the promoter regions of genes central to pluripotency maintenance as well as genes involved in early lineage specification (Figure 6H, I, Supplementary 6H–K). These shifts in chromatin state suggest a broad reprogramming of transcriptional control networks that likely underlie the impaired differentiation phenotype observed in AMDHD2-deficient mESCs (Figure 4A–D, G–J). Furthermore, the acetylation changes were accompanied by alterations in repressive histone modifications, as indicated by a global reduction in H3K27me3 levels in the KO line (Supplementary Figure 6K–O), consistent with a disturbed balance between activating and repressive histone marks. Taken together, these findings support a model in which AMDHD2 loss perturbs intracellular acetate pools, leading to widespread changes in epigenetic regulation that disrupt the transcriptional dynamics required for proper differentiation.

Finally, to determine whether the loss of AMDHD2 function affects differentiation in a conserved manner, we examined the impact of AMDHD2 KD on the differentiation of human induced pluripotent stem cells (hiPSCs). We treated the cells with either a non-targeted control siRNA or an AMDHD2-specific siRNA, followed by a seven-day differentiation protocol. Knockdown efficiency of AMDHD2 was confirmed by WB analysis (Figure 6J) and qPCR (Supplementary Figure 6P). We were able to measure increases in the expression levels of all lineage markers throughout the differentiation process (Figure 6K–N), indicating a successful heterogenous differentiation. However, we observed a reduction in the expression levels of most lineage markers tested on day 7 in AMDHD2 siRNA-treated cells compared to control cells (BRACHYURY: primitive streak, MSX1: mesoderm, OTX2: neuroectoderm, GATA6: endoderm; Figure 6J–M). These data highlight the importance of AMDHD2 activity in the productive progression of differentiation across species.

## Discussion

The hexosamine biosynthetic pathway (HBP) integrates multiple critical intermediates from various metabolic pathways for the synthesis of UDP-GlcNAc, a key metabolite with established roles in regulating SC fate and identity^23–25, 28,44^. In this study, we identified AMDHD2 as a novel regulator of the cyto-nuclear acetate/acetyl-CoA pool through its function in deacetylating the HBP intermediate GlcNAc6P supplied by *de novo* synthesis and the GlcNAc salvage pathway (Figure 7).

**Figure 7:**
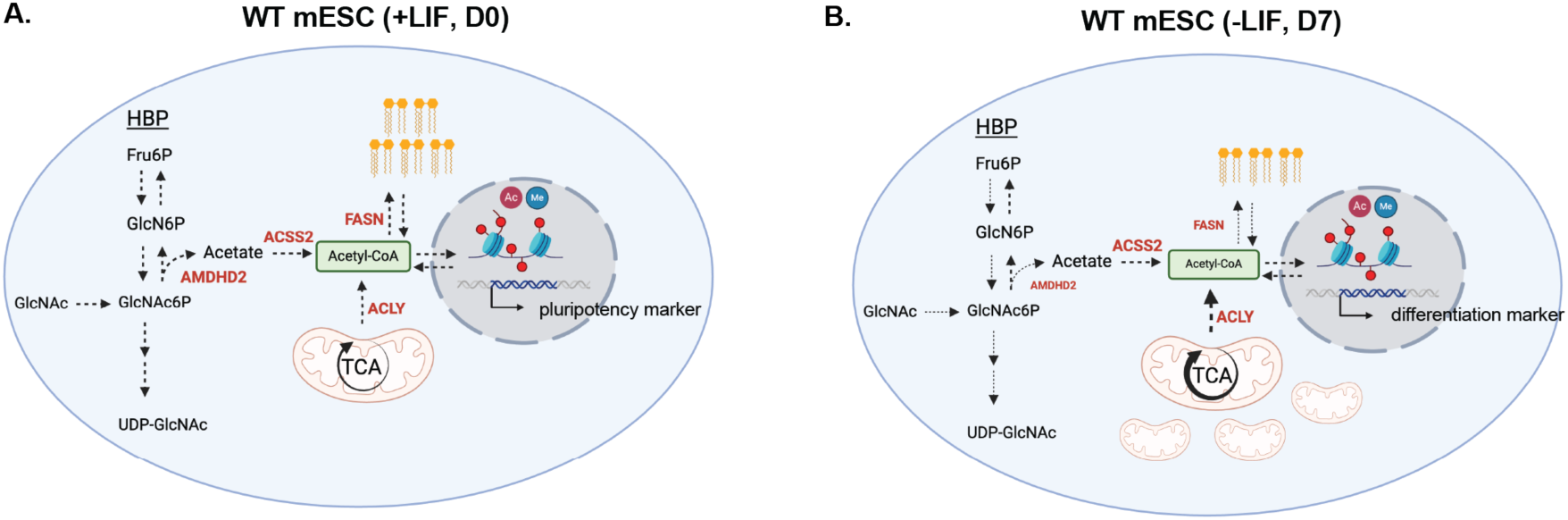
AMDHD2 mediates a dual function of the hexosamine biosynthetic pathway by regulating acetyl-CoA homeostasis in mESCs. **A.** In pluripotent mESCs (+LIF, D0) the hexosamine biosynthetic pathway (HBP) generates the end product UDP-GlcNAc by using a combination of *de novo* synthesis and the GlcNAc salvage pathway. The intermediate GlcNAc6P is steadily deacetylated by AMDHD2, releasing acetate which is further converted to acetyl-CoA by ACSS2. The generated cytosolic acetyl-CoA pool is used for lipid synthesis and histone modifications, which regulate the expression of genes important for pluripotency. **B.** After 7 days of mESC differentiation induced by LIF deprivation (-LIF, D7), glucose flux through *de novo* synthesis and GlcNAc salvage flux into UDP-GlcNAc are reduced. AMDHD2 expression and likely its activity is reduced, thereby generating less acetate. Upon differentiation glycolysis is reduced, while the TCA cycle activity increases, elevating cytosolic acetyl-CoA supply via ACLY. FASN (and lipid synthesis) is downregulated. Overall, reduced cytosolic acetyl-CoA levels alter the epigenetic landscape, thereby regulating gene expression of transcripts involved in differentiation and development.

### N-acetyl-Glucosamine-6-phosphate is a critical metabolite in stem cell differentiation

Our findings reveal a dual role for the HBP in SC differentiation: in addition to its known role in O-GlcNAcylation of critical pluripotency markers, we demonstrate that the intermediate GlcNAc6P plays a crucial role in differentiation^23–26, 45, 46^. To maintain GlcNAc6P levels, pluripotent cells use a combination of *de novo* synthesis and the GlcNAc salvage pathway (Figure 2F, K). GlcNAc6P is then converted to GlcN6P by AMDHD2 (Figure 3B), releasing acetate (Figure 5A), which ACSS2 subsequently uses to generate acetyl-CoA (Figure 5F). This pathway represents an additional mechanism for maintaining the nucleo-cytosolic acetyl-CoA pool in pluripotent stem cells, complementing classically studied sources such as mitochondrial-derived citrate^39, 47^. Relatedly, pluripotent ESCs exhibit low TCA cycle activity, which increases markedly upon differentiation^48^. Therefore, while mitochondrial-derived citrate is exported to the cytosol and converted by ACLY as a major source of acetyl-CoA in differentiated cells, acetate conversion via ACSS2 may play a particularly important role in ESCs. Our model supports this, as we observed increased expression of TCA cycle enzymes, metabolite levels and glucose flux into the TCA cycle during differentiation (Figure 1H, J).

Loss of AMDHD2 function resulted in impaired differentiation (Figure 4A– D), emphasizing its importance in early cell fate transitions. To validate this mechanism, we phenocopied the impaired differentiation observed in AMDHD2 KO cells by conditionally depleting AMDHD2 (via degron-mediated depletion and siRNA-mediated knockdown) in both mESCs and hiPSCs (Figure 4G–J, Figure 6K–N). Additionally, ACSS2 inhibition resulted in a similar differentiation defect (Figure 5H–K, Supplementary Figure 5F–I). These findings highlight the critical role of AMDHD2-mediated acetate release in early differentiation and SC identity transitions.

### AMDHD2 modulates the production of acetate in pluripotent stem cells

Acetyl-CoA is a central metabolite that links metabolism to epigenetic regulation by driving histone acetylation, thereby influencing gene expression and cell fate decisions^39, 49^. During the first 24–48 hours of differentiation, metabolic rewiring leads to decreased histone acetylation, a necessary step to exit pluripotency^7^. Our results show that dysregulation of acetate levels by AMDHD2 depletion during the first 72 h of differentiation is sufficient to impair long-term differentiation outcomes (Figure 4G–J), highlighting its essential function in early developmental transitions. In line with this, we measured a reduction in GlcNAc-derived acetate levels in AMDHD2 KO and steady-state acetate levels upon AMDHD2 depletion. While acetate supplementation could rescue the depletion of AMDHD2 completely (Figure 5B–D) and re-expression of AMDHD2 provided a partial rescue (Supplementary Figure 5C–E), excessive amounts of acetate impaired differentiation (Figure 5B–D), in line with previous publications ^50^. In addition, we could show that GlcNAc significantly contributes to acetyl-CoA levels in WT cells (Figure 5F) and we measured reduced GlcNAc-derived acetyl-carnitine in AMDHD2 KO cells as a proxy for acetyl-CoA levels (Figure 6A), suggesting that AMDHD2 can indirectly limit acetyl-CoA levels.

It has been shown that cells actively maintain acetyl-CoA homeostasis through several mechanisms. For example, ACLY depletion leads to increased ACSS2 expression in cancer cells and mouse embryonic fibroblasts^9–11^. Similarly, we observed compensatory mechanisms in AMDHD2 KO cells, including upregulation of ACSS2 and downregulation of FASN (Figure 4K, L), further highlighting the importance of this tightly regulated metabolic network in maintaining acetyl-CoA levels.

### Subcellular Metabolite Regulation and Future Directions

Emerging evidence highlights the significance of subcellular metabolite compartmentalization. Ma et al. demonstrated that effector CD8+ T cells preferentially utilize acetate and ACSS2 in complex with p300 to regulate histone acetylation, whereas exhausted T cells rely on citrate and ACLY in a complex with KAT2A, leading to distinct histone modification patterns influencing cell fate^51^. Similarly, ACSS2 has been implicated in lactyl-CoA synthesis, interacting with LDHA and KAT2A to promote histone lactylation and gene regulation in glioblastoma cells^52^. These findings suggest broader epigenetic roles for ACSS2 beyond acetyl-CoA generation, raising intriguing possibilities for AMDHD2 and ACSS2 interactions in SCs. Future studies should explore whether AMDHD2 and ACSS2 colocalize in the nucleus and cooperatively regulate histone modifications to influence SC fate decisions.

## Conclusion

Maintaining the acetyl-CoA pool in a SC-specific context is a complex, finely tuned process. Our study introduces a novel regulatory layer by identifying AMDHD2 as a supplier of intracellular acetate. The steady conversion of GlcNAc6P to GlcN6P by AMDHD2 releases acetate which is particularly required in pluripotent SCs. Differentiation leads to reduced AMDHD2 expression and decreased pathway activity (Figure 2H, M; Supplementary Figure 2A, B, Figure 3K, L), which aligns well with an increased reliance on the TCA cycle for acetyl-CoA supply. By demonstrating an HBP-mediated mechanism in addition to UDP-GlcNAc production, we provide new insights into how metabolic pathways interface with epigenetic regulation to control cell fate transitions (Figure 7).

---

**Supplementary Figure 1.**
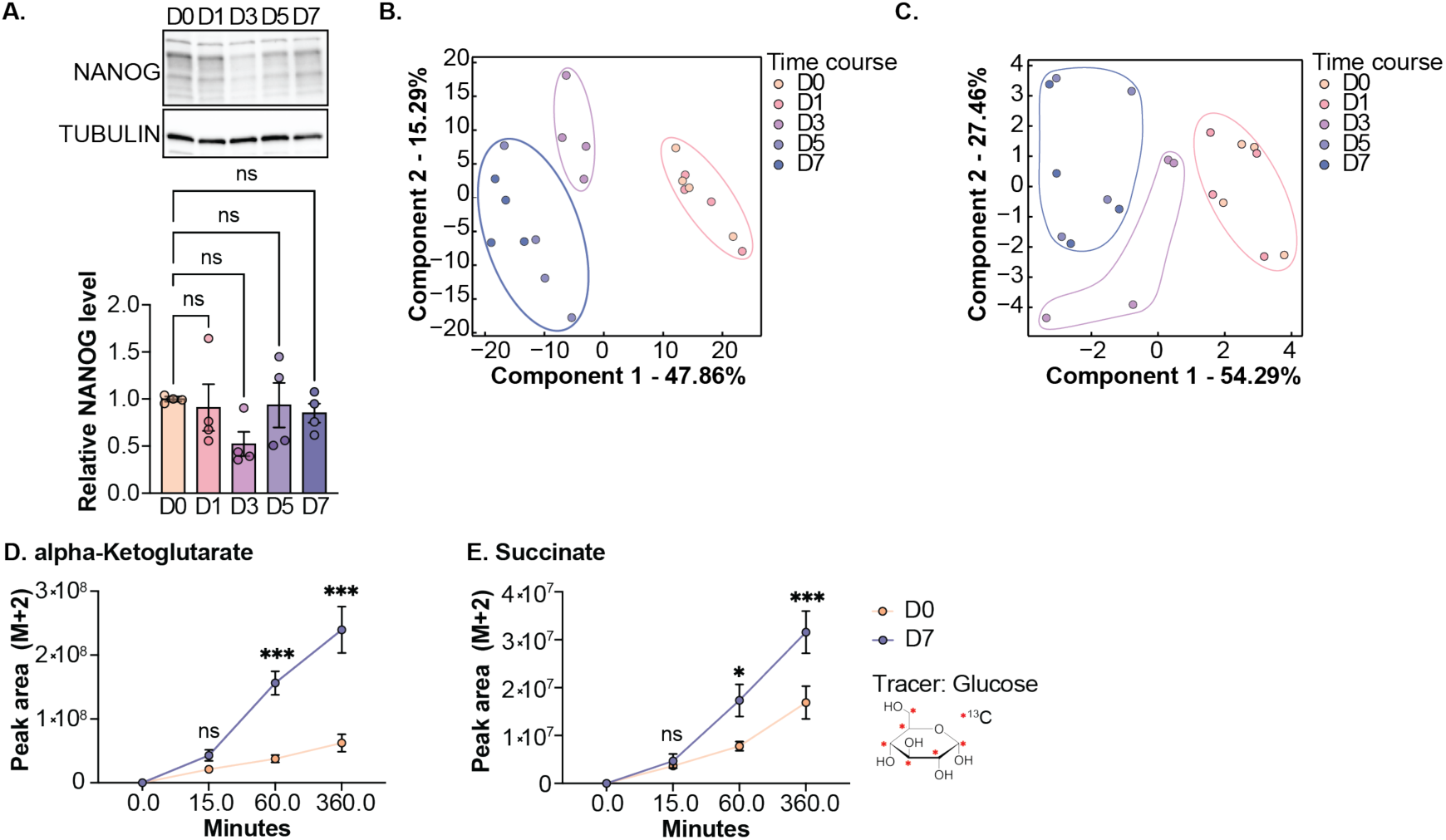
**A.** Western blot analysis and quantification of NANOG in WT cells during differentiation (mean ± SEM, n=4, One-way ANOVA Dunnett post-test). **B.** Principal component analysis (PCA) of proteomics dataset of WT mESCs during the differentiation process. **C.** PCA of metabolomics dataset of WT mESCs during the differentiation process. **D.** Pluripotent and 7-day differentiated (D7) WT cells were labelled with ^13^C_6_-glucose. Shown is the labelled fraction (M+2) of alpha-ketoglutarate at 0, 15, 60, and 360 minutes of labelling (mean ± SEM, n=4, * p<0.05, *** p<0.001, Two-way ANOVA Šídák’s post-test). **E.** Same setup as in E. for succinate.

**Supplementary Figure 2.**
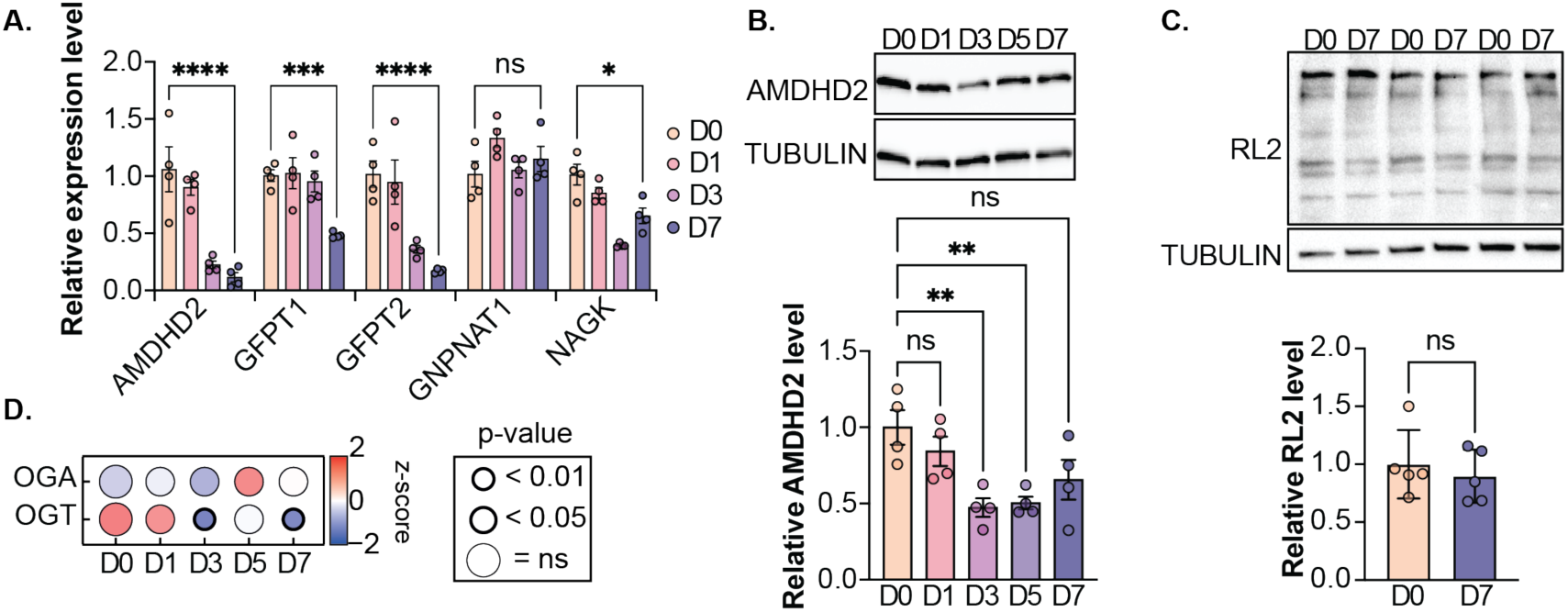
**A.** Relative mRNA levels of the HBP enzymes in WT cells during the differentiation process (mean ± SEM, n=4, * p<0.05, *** p<0.001, **** p<0.001 Two-way ANOVA Tukey post-test). **B.** Western blot analysis and quantification of AMDHD2 in WT cells during differentiation (mean ± SEM, n=4, ** p<0.01, One-way ANOVA Tukey post-test). **C.** Western blot analysis and quantification of O-GlcNAcylated proteins using the RL2 antibody in WT cells during differentiation (mean ± SD, n=5, unpaired, one-tailed t-test). **D.** Proteomics data for OGA and OGT in WT cells during differentiation. Significances are indicated by different circle size and thickness (n=4).

**Supplementary Figure 3.**
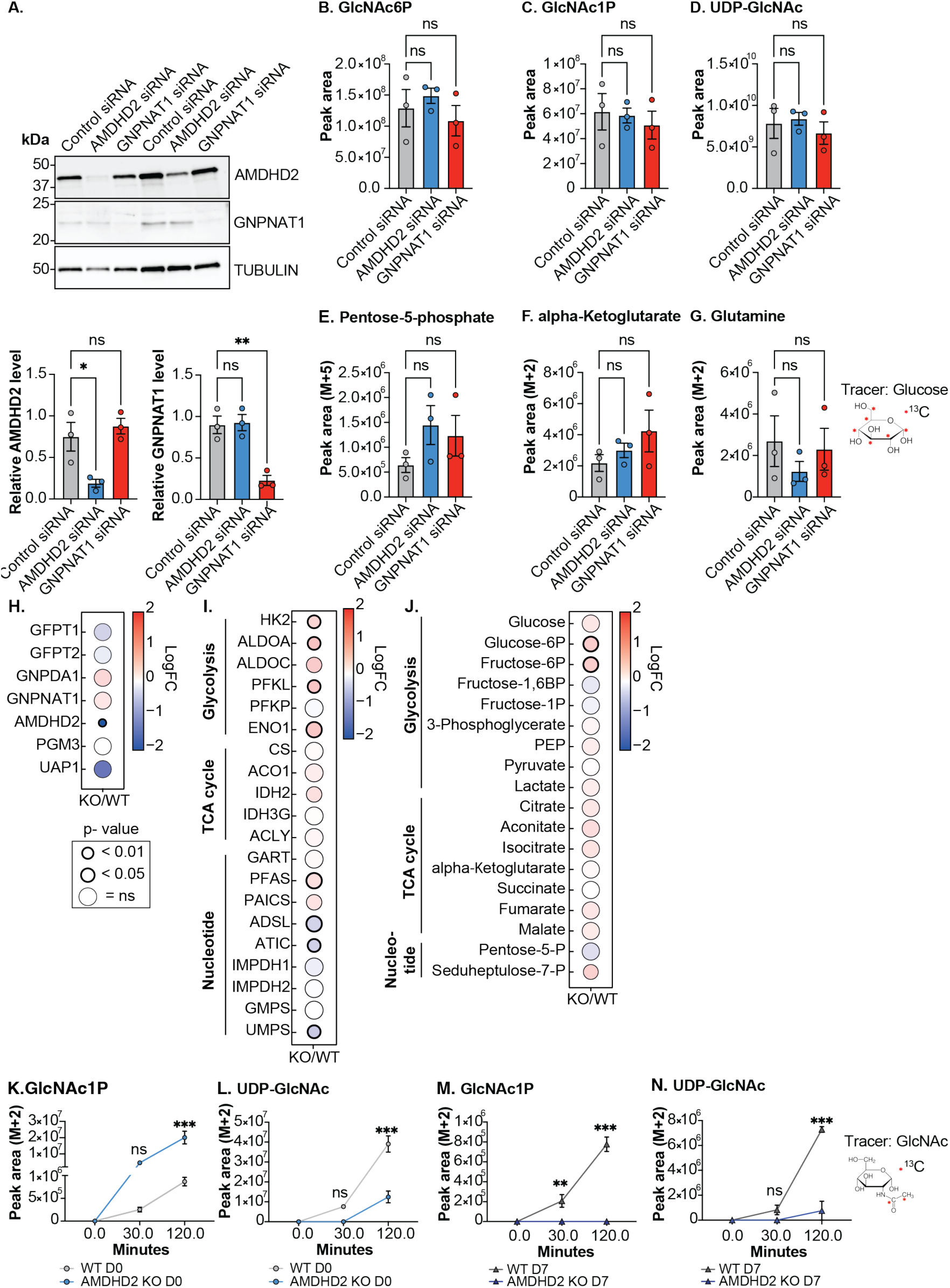
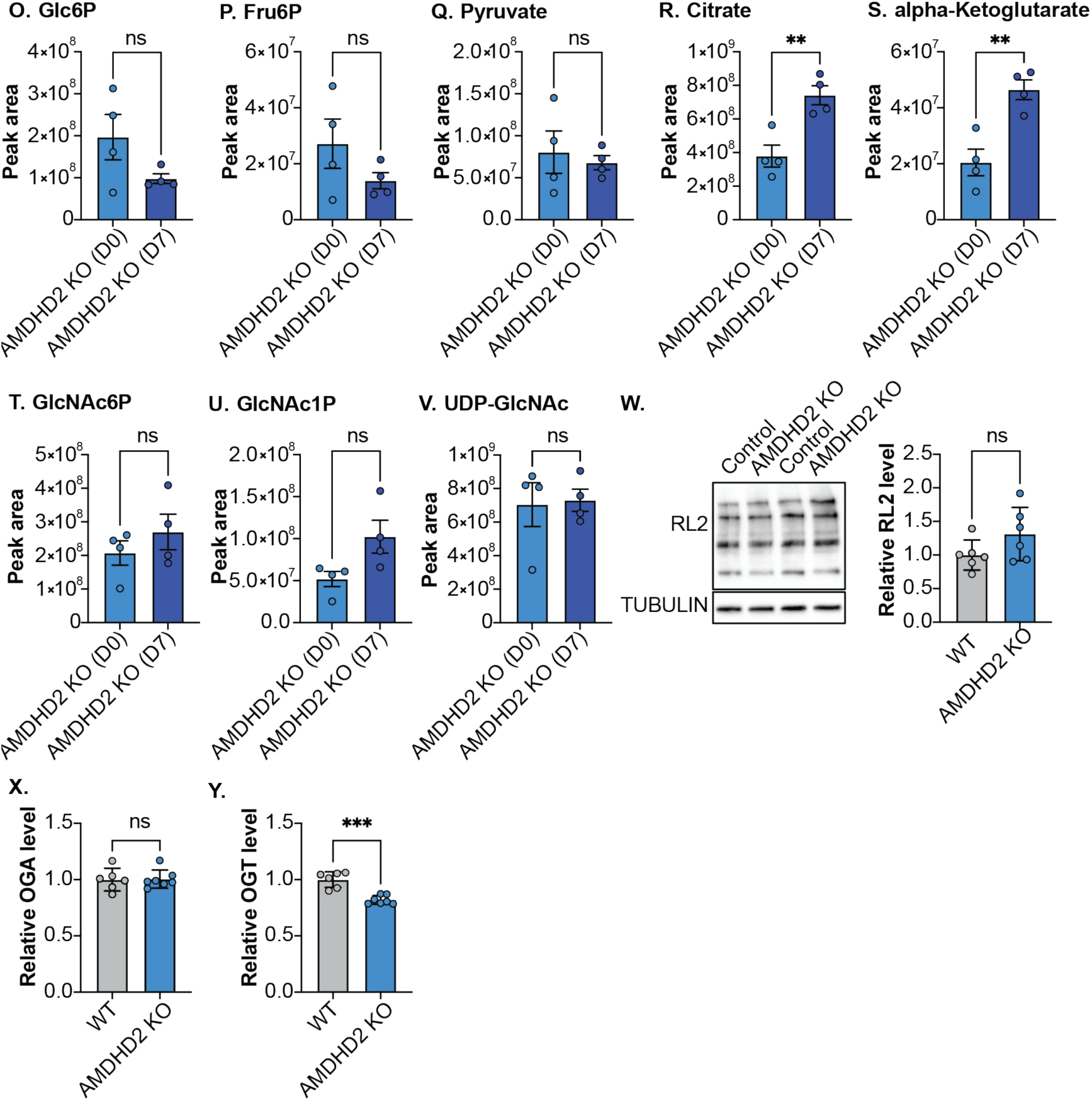
**A.** Western blot analysis and quantification of AMDHD2 and GNPNAT1 in WT cells 48 h post transfection with non-targeting siRNA or siRNA against AMDHD2 or GNPNAT1, respectively (mean ± SD, n=3, * p<0.05, ** p<0.01, One-way ANOVA Dunnett post-test). **B.** Same setup as in A. After 48 h, cells were collected for metabolomics analysis and steady-state GlcNAc6P levels were measured (mean ± SEM, n=4, One-way ANOVA Tukey post-test). **C.** Same setup as in B. for GlcNac1P, as in **D.** for UDP-GlcNAc. **E.** Same setup as in A. After 48 h, cells were labelled with ^13^C_6_-glucose. Depicted is the labelled fraction (M+5) of pentose-5-phosphate after 60 min of labelling (mean ± SEM, n=4, One-way ANOVA Tukey post-test). **F.** Same setup as in E for the M+2 fraction of alpha-Ketoglutarate. **G.** Same setup as in F. for glutamine. **H.** Proteomics data of the HBP enzymes in WT compared to AMDHD2 KO cells. Significances are indicated by differences in circle size and thickness (n=6). **I.** Same setup as in H. for other metabolic enzymes. **J.** Metabolomics data of WT cells compared to AMDHD2 KO cells. Significances are indicated by differences in circle size and thickness (n=6). **K.** Pluripotent WT and AMDHD2 KO mESCs were labelled with ^13^C_2_-GlcNAc. Depicted is the labelled fraction (M+2) of GlcNAc1P, after 0, 30 and 120 min of labelling (mean ± SEM, n=4, *** p<0.001, Two-way ANOVA Šídák’s post-test). **L.** Same setup as in K for UDP-GlcNAc. **M.** D7 differentiated WT and AMDHD2 KO mESCs were labelled with ^13^C_2_-GlcNAc. Depicted is the labelled fraction (M+2) of GlcNAc1P after 0, 30 and 120 min of labelling (mean ± SEM, n=4, *** p<0.001, Two-way ANOVA Šídák’s post-test). **N.** Same setup as in M. for UDP-GlcNAc. **O.–V.** IC-MS analysis of glycolytic metabolites (mean ± SEM, n=4, ** p<0.01, unpaired, one-tailed t-test).: **O.** Glucose-6-phosphate (Glc6P), **P.** Fructose-6-phosphate (Fru6p), **Q.** Pyruvate, **R.** Citrate, **S.** alpha-Ketoglutarate, **T.** GlcNAc6P, **U.** GlcNAc1P. **V.** UDP-GlcNAc. **W.** Western blot analysis and quantification of O-GlcNAcylated proteins using the RL2 antibody in WT and AMDHD2 KO cells (mean ± SD, n=6, unpaired-test). **X.** Proteomics data of OGA in WT and AMDHD2 KO mESCs (mean ± SD, n=6, *** p<0.001, unpaired, one-tailed t-test). **Y.** Same setup as in X. for OGT.

**Supplementary Figure 4.**
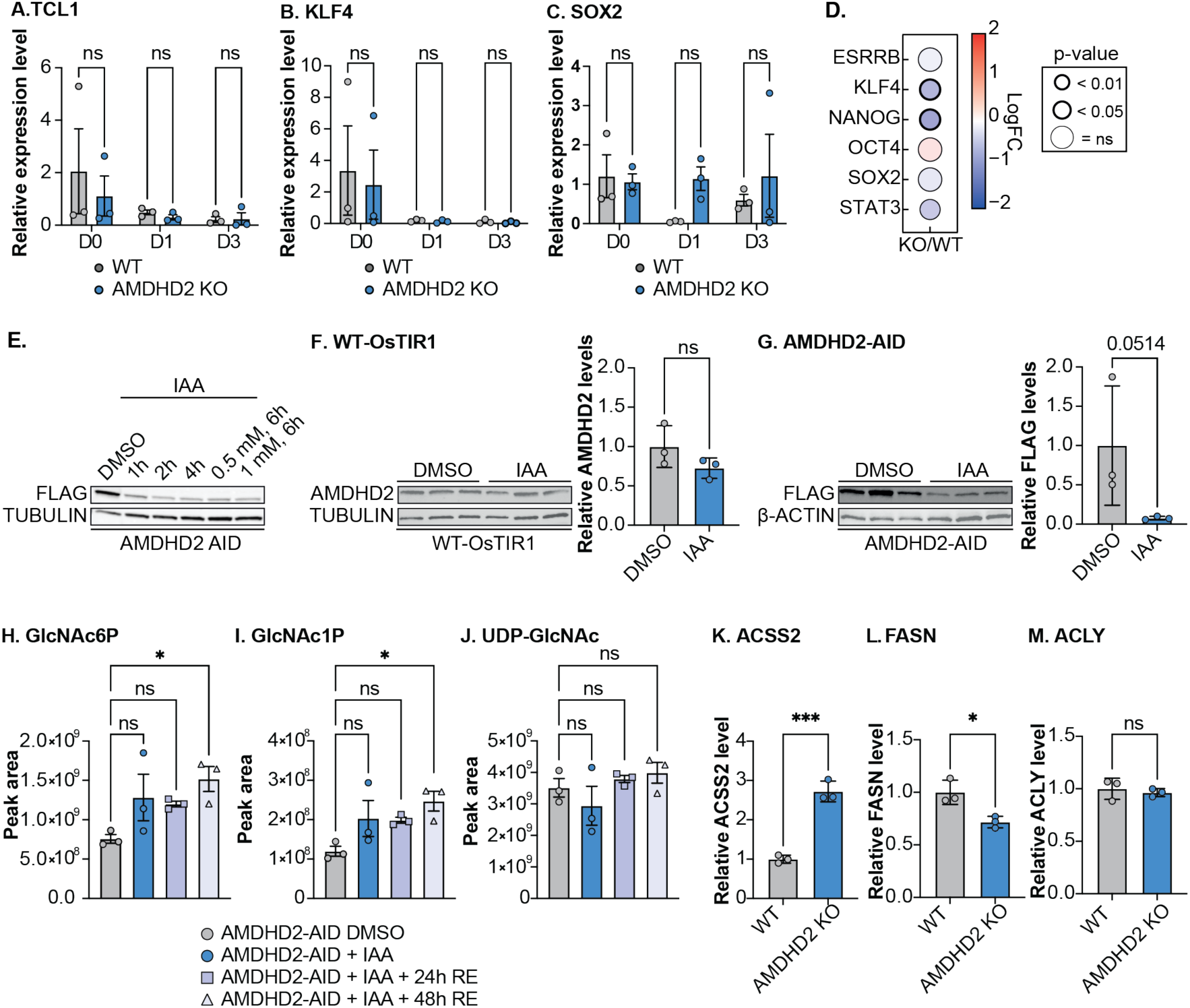
**A.** Relative mRNA levels of TCL1 in WT and AMDHD2 KO cells during early differentiation (mean ± SEM, n=3, One-way ANOVA Tukey post-test). **B.** Same setup as in A. for KLF4, in **C.** for SOX2. **D.** Proteomics data of pluripotency markers in WT and AMDHD2 KO cells. Significances are indicated by differences in circle size and thickness (n=6). **E.** Western blot analysis to confirm functionality of the AMDHD2-AID system. AMDHD2-FLAG-AID cells co-expressing the OsTIR1-V5 ligase were treated with DMSO or IAA for 1, 2, 4, 6 h with 0.5 mM IAA or 6 h with 1 mM IAA. **F.** Western blot analysis and quantification to confirm functionality of the AMDHD2-AID system. WT cells only expressing the OsTIR1-V5 ligase were treated with DMSO or 0.5 mM IAA 6 h (mean ± SD, n=3, unpaired, one-tailed t-test). **G.** Same setup as in F for AMDHD2-AID cells co-expressing the OsTIR1-V5 ligase. **H.** IC-MS analysis of the HBP intermediates in AMDHD2-FLAG-AID cells treated with DMSO or 0.5 mM IAA for 6 h, followed by 24 h or 48 h recovery (mean ± SEM, n=3, * p<0.05, One-way ANOVA Dunnett post-test). **I.** Same setup as in H. for GlcNac1P, in **J.** for UDP-GlcNAc. **K.–M.** Proteomics data of **K.** ACSS2, **L.** FASN and **M.** ACLY in WT and AMDHD2 KO cells (mean ± SD, n=3, * p<0.05, *** p<0.001, unpaired, one-tailed t-test).

**Supplementary Figure 5.**
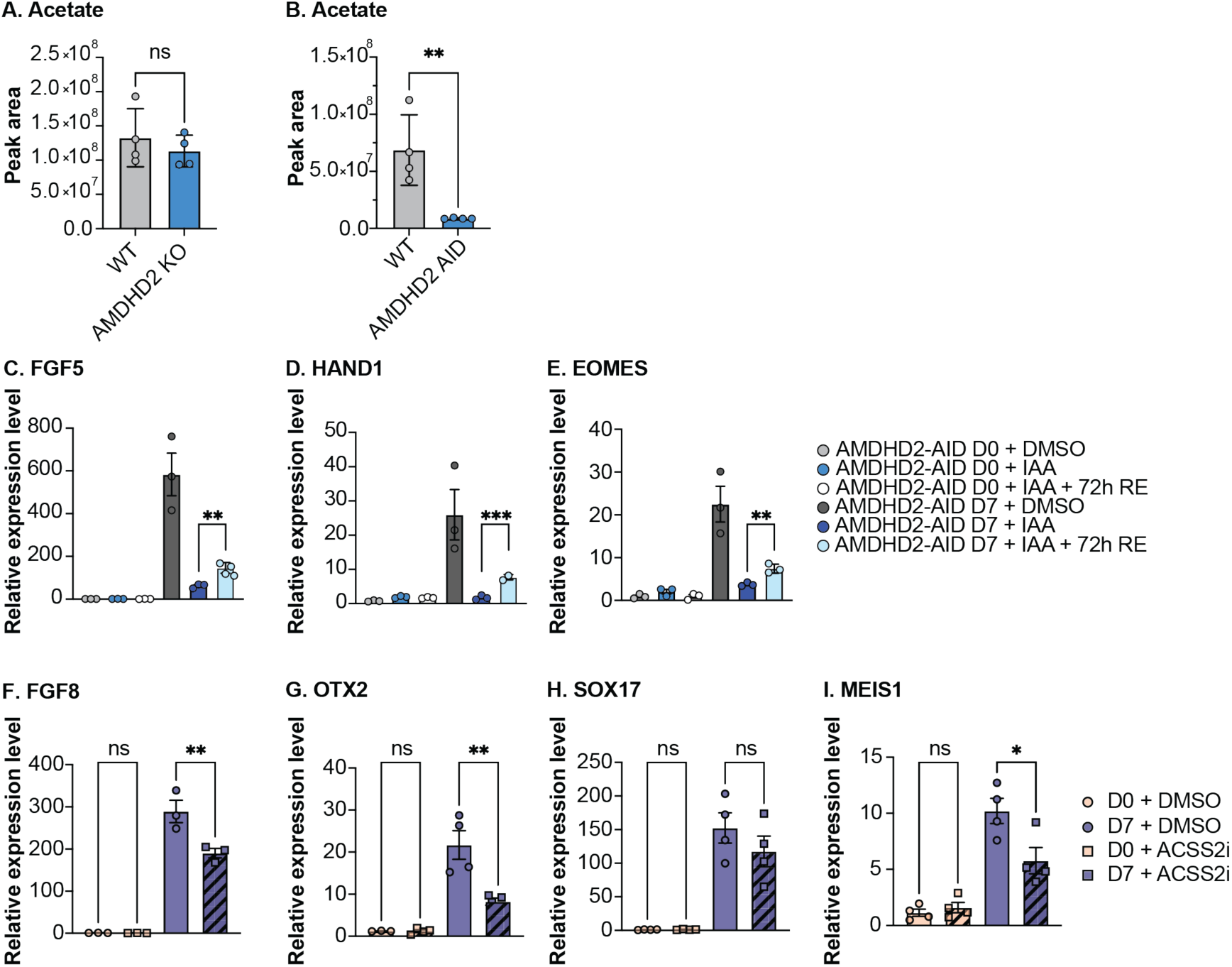
**A.** UPLC-HRMS analysis of acetate in WT and AMDHD2 KO cells after 48 hours of ACSS2 inhibition (10 µM; mean ± SEM, n=4). **B.** UPLC-HRMS analysis of acetate in WT and AMDHD2 AID cells after 6 hours of IAA and ACSS2 inhibitor treatment (10 µM, mean ± SEM, n=4, ** p<0.01). **C.** Relative mRNA expression of FGF5 in pluripotent and D7 AMDHD2-AID cells. All cells were either treated with DMSO or IAA for 6 hours before differentiation (D0). In addition, cells were left to recuperate (RE) for 72 hours or used for differentiation without recovery of AMDHD2 levels (mean ± SEM, n=3, ** p<0.01, *** p<0.001, unpaired, one-tailed t-test). **D.** Same setup as in C. for HAND1, in **E.** for EOMES. **F.** Relative mRNA expression of FGF8 in pluripotent and D7 WT cells. Cells were treated with DMSO or 1 µM ACSS2 inhibitor for 24 hours at D0 or with DMSO or 0.1 µM ACSS2 inhibitor for the first 72 h of differentiation (mean ± SEM, n=4, * p<0.05, ** p<0.01, One-way ANOVA Tukey post-test). **G.** Same setup as in F. for OTX2, in **H.** for SOX17, and **I.** for MEIS1.

**Supplementary Figure 6.**
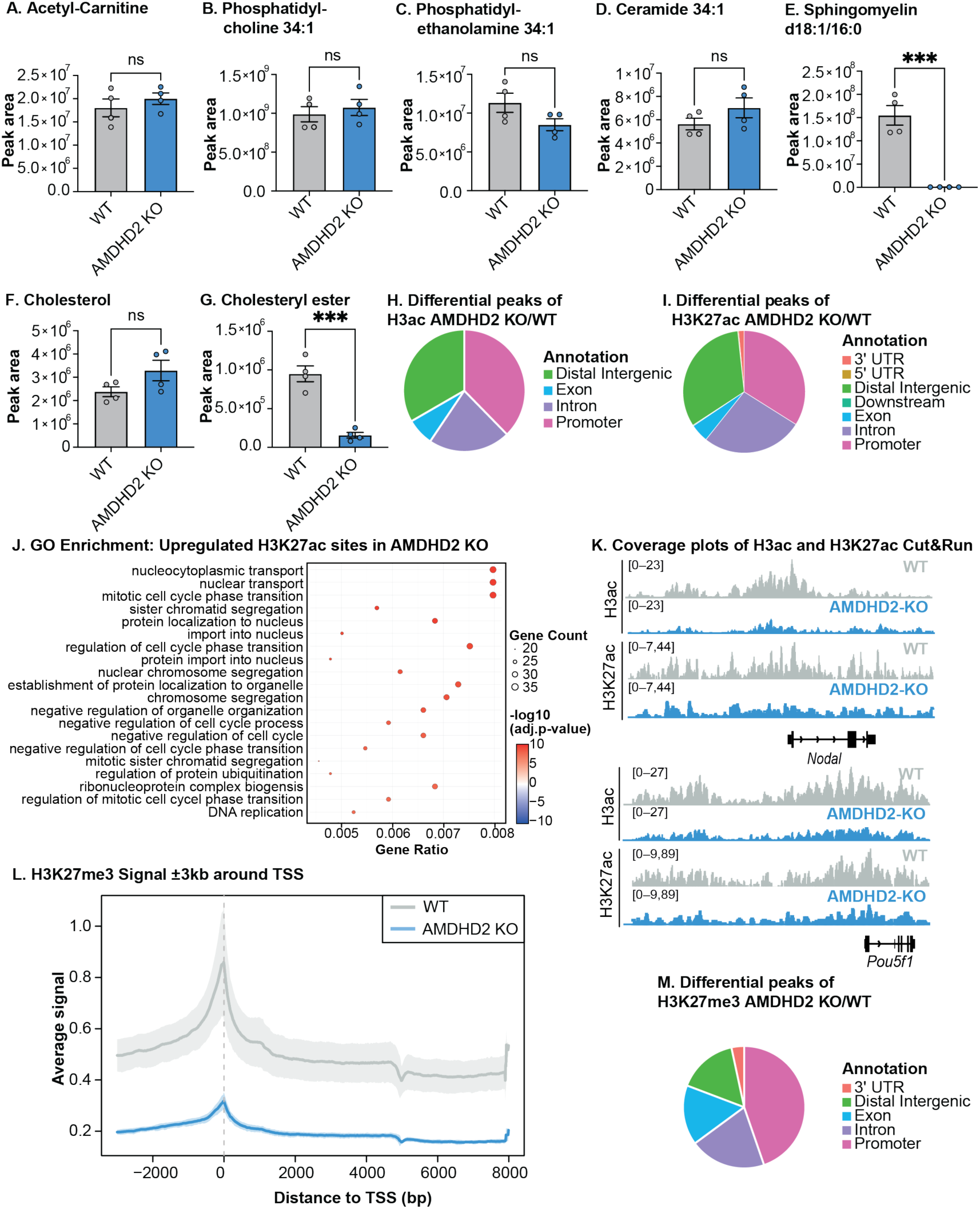
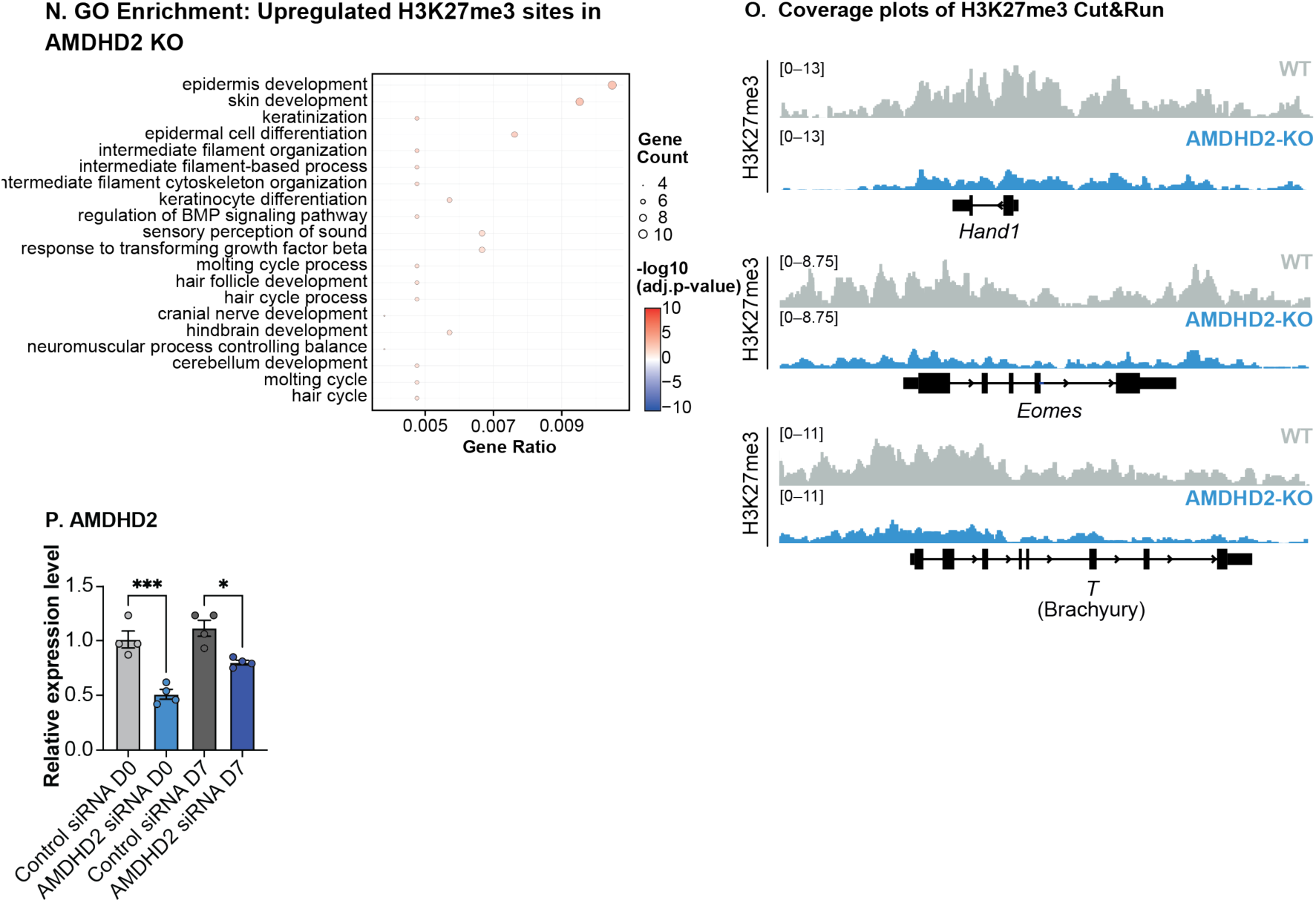
**A.** Measurement of acetyl-carnitine pluripotent WT and AMDHD2 KO cells (mean ± SEM, n=4, *** p<0.001, unpaired, one-tailed t-test). **B.** Same setup as in A. for phosphatidylcholine, in **C.** for phosphatidylethanolamine, in **D.** for ceramide and in **E.** for sphingomyelin, in **F.** for cholesterol and for **G.** in cholesteryl ester. **H.** Differential peaks of H3ac CUT&RUN peaks comparing data from AMDHD2 KO to WT cells. **I.** Same setup as in H. for H3K27ac CUT&RUN data. **J.** GO enrichment of the genes with upregulated H3K27ac signal in AMDHD2 KO cells. **K.** Coverage plots of H3ac and H3K27ac CUT&RUN data for WT and AMDHD2 KO cells surrounding the genes *Nodal* and *Pou5f1*. **L.** Mapping of H3K27-methylation (H3K27me) CUT&RUN signal 3 kb upstream and downstream of the transcription start site from WT and AMDHD2 KO cells. **M.** Differential peaks of H3K27me CUT&RUN peaks comparing data from AMDHD2 KO to WT cells. **N.** GO enrichment of the genes with upregulated H3K27me signal in AMDHD2 KO cells. **O.** Coverage plots of H3ac and H3K27ac CUT&RUN data for WT and AMDHD2 KO cells surrounding the genes *Hand1*, *Eomes* and *T* (BRACHYURY). **P.** Relative mRNA expression of AMDHD2 in pluripotent human iPSCs and 7-day differentiated human iPSCs following treatment with control siRNA or siRNA AMDHD2 (mean ± SEM, n=4, * p<0.05, One-way ANOVA Tukey post-test).

## Methods

### Cell lines and culture conditions

AN3-12 mouse embryonic stem cells were cultured as previously described (Elling et al., 2011). In brief, DMEM high glucose (Thermo Fisher Scientific) was supplemented with L-glutamine, fetal bovine serum (15%), penicillin/streptomycin, non-essential amino acids, sodium pyruvate (all Thermo Fisher Scientific), β-mercaptoethanol and LIF (EMBL) and used to culture cells at 37°C in 5% CO_2_ on non-coated tissue culture plates. Seeding numbers for 10 cm dishes ranged between 180,000–350,000 cells and for 6 cm dishes from 50,000–100,000 cells.

For induction of spontaneous differentiation of AN3-12 cells, cells were counted with a TC20 automated cell counter (BioRad) and seeded in a 6-well plate and incubated for one, three, five or seven days in medium without LIF with a regular exchange of medium (for D0: 400,000 cells, for D1: 100,000 cells, for D3: 20,000 cells, for D7: 5,000 cells). AMDHD2 KO AN3-12 cells had been previously generated in Kroef et al., 2022. Mycoplasma testing was performed at regular intervals and always tested negative.

iPSCs (UKBi017-A, https://hpscreg.eu) were cultured in feeder-free conditions in StemMACS iPS-Brew (Miltenyi Biotec) on vitronectin-coated 6-well plates. The medium was changed daily, and PBS-EDTA was used for passaging. For undirected differentiation experiments, iPSC colonies were first dissociated into single cells using accutase. Approximately 1 million cells were taken up in 2 mL StemMACs iPS-Brew medium, supplemented with 10 μM ROCK inhibitor (Y-27632, Miltenyi Biotec). iPSCs were then grown on non-coated 6-well plates for 5 days to allow the formation of embryoid bodies (EBs). Thereafter, EBs were transferred onto gelatin-coated plates and the medium was switched to serum-containing MEF-medium (DMEM, 10 % FBS, 1 % nonessential amino acids, 2 mM of L-glutamine, 0.1 mM of β-mercaptoethanol), which induces spontaneous differentiation. The medium was replaced every other day and cells were collected for biochemical analysis at the indicated timepoints.

### Cell sorting

To maintain a haploid or diploid cell population, cells were stained with 10 μg/ml Hoechst 33342 (Thermo Fisher Scientific) for 30 min at 37°C. To exclude dead cells, propidium iodide (Sigma-Aldrich) staining was added. Cells were sorted for DNA content on a FACSAria Fusion sorter and flow profiles were recorded with the FACSDiva software (BD Franklin Lakes). Haploid cells were used solely for cloning procedures, while all other experiments were conducted using AN3-12 cells in the diploid state.

### Generation of cells expressing the cytosolic acetyl-CoA sensor

For introducing the acetyl-CoA sensor encoding gene into the *Rosa26* locus, we used the protocol and constructs as previously described ^40, 53^. The acetyl-CoA sensor encoding sequence was amplified from pLJM1-Cytoplasmic PancACe (RRID: Addgene_215708) by PCR and introduced in to the donor plasmid for knock-in using restriction-free cloning. The CMV promoter was further exchanged for an EF1α promoter. Haploid mESCs were transfected with 1 µg of Cas9-mRFP encoding plasmid and 3 ug of the donor plasmid using Lipofectamine 3000 (Thermo Fisher Scientific) according to the manufacturer’s protocol. GFP and mRFP positive cells were sorted by the BD FACSAria Cell sorter and cultured for several days. After single cell sorting into 96 well plate, clones were analyzed for proper transgene insertion by genotyping PCR and by measuring the intensity of the acetyl-CoA sensor by flow cytometry (BD FACSCantoII).

### Flow cytometry for PancACe sensor-expressing cells after glucose and glutamine deprivation and refeeding

The cells were cultured in regular medium or in glucose and glutamine free DMEM (Thermo Fisher Scientific) supplemented with 15 % fetal bovine serum, penicillin/streptomycin, non-essential amino acids, sodium pyruvate, β-mercaptoethanol and LIF for 4 h to induce starvation. Afterwards, the cells were incubated for 1 h with either regular medium, medium supplemented with 10 mM GlcNAc as the sole carbon source, or medium containing 10 mM GlcNAc along with 10 µM ACSS2 inhibitor (SelleckChemicals, S8588). Harvested cells were resuspended in glucose and glutamine-free FACS buffer co supplemented with 15 % fetal bovine serum, 1xHBSS, penicillin/streptomycin, non-essential amino acids, sodium pyruvate, β-mercaptoethanol and LIF and analyzed by FACS (BD FACSCantoII). Dead cells were excluded by DRAQ7 (BioLegend) staining. The data were analyzed using the FlowJo software (v.10.10).

### Generation of AMDHD2-FLAG-AID cells

For introducing the OsTIR1 gene into the TIGRE locus, we used the protocol and constructs as previously described^38^. To tag endogenous AMDHD2 at its C-terminus with an AID–FLAG, we generated a targeting construct with a neomycin resistance, based on plasmids which were kindly provided by F. Dossin (formerly Group of Edith Heard). 500-bp homology arms upstream and downstream of the coding sequence (but excluding the stop codon) were amplified from mouse genomic DNA by PCR and introduced in to the respective plasmid using restriction-free cloning. DNA template sequences for small guide RNAs targeting AMDHD2 were designed online (http://crispor.org, Supplementary Table 2), purchased from Sigma, and cloned into the Cas9-GFP expressing plasmid PX458 (RRID: Addgene_48138). Haploid mESCs were transfected with 4 µg DNA using Lipofectamine 2000 (Life Technologie) according to the manufacturer’s protocol. The selection was performed with 0.6 mg/ml G418 (Gibco) for several days. Afterwards single clones were analyzed for proper transgene insertion by genotyping PCR before being sequenced to exclude potential mutations.

### RNA isolation and qPCR

RNA was isolated using the Quick RNA Miniprep kit (Zymogen), as recommended by the manufacturer. 1.0 μg total RNA was used to synthesize cDNA using the iScript cDNA Synthesis kit (BioRad). RT-qPCR was performed using the 2x Universal SYBR-Green Fast qPCR mix on a CFX384 Touch Real-Time PCR Detection System. Gene expression values were normalized using RPL37A and are shown as a relative fold change to the value of control samples. All experiments were performed in biological triplicates and error bars indicate ± standard deviation as assayed by the ΔΔCt method. All primers are listed in Supplementary Table 1.

### siRNA-mediated knockdown

siRNA-mediated knockdown was carried out to reduce the expression of target genes. Cells were transfected with 50 nM of either specific siRNA or non-targeting control siRNA (mouse: L-064305-01-0005, control: D-001810-10-05; Dharmacon, Lafayette, CO) using Lipofectamine RNAiMAX (Invitrogen) following the manufacturer’s protocol. In brief, siRNA and Lipofectamine RNAiMAX were each diluted in Opti-MEM medium (Gibco), mixed, and incubated at room temperature for 20 minutes to form complexes. The resulting siRNA-lipid complexes were then added dropwise to cells at 60–70% confluency. mESCs were incubated for 48 hours post transfection to allow for efficient knockdown before conducting further experiments. Knockdown efficiency was assessed by Western blot analysis.

iPSCs were transfected with synthetic siRNAs (human: L-008651-00-0005; Dharmacon) using Lipofectamine RNAiMax (Thermo Fisher Scientific) according to the manufacturer’s instructions. Briefly, iPSC colonies were first dissociated into single cells using accutase. Approximately 1 million cells were then incubated in 100 µL of the respective transfection mix for 10 min before continuing with the differentiation experiments.

### Metabolite extraction of polar metabolites

A wash buffer solution was prepared by dissolving 0.7207 g of ammonium carbonate (Sigma-Aldrich) in 100 ml of LC-MS ultra-grade water. Alternatively, a 750 mM stock solution of ammonium carbonate (7.207 g per 100 mL) was prepared in LC-MS ultra-grade water and diluted 1:10 (v/v) with the same water prior to use. Glacial acetic acid (Sigma-Aldrich) was added at a concentration of 2 µL per mL of wash buffer (i.e. 200 µL per 100 mL). The pH was then adjusted in 10 µL increments until a final pH of 7.4 ± 0.02 was achieved. The wash buffer was stored at 4 °C until use. For buffers stored for more than 1–2 days, the pH was rechecked and readjusted with small volumes of glacial acetic acid as required. The extraction buffer was freshly prepared prior to use by mixing LC-MS grade methanol, acetonitrile and water at a ratio of 40:40:20 (v:v:v). Internal standards were added to the buffer as follows (per 100 mL total volume): 10 µL of 2.5 mM U-^13^C^15^N-labelled amino acids; 10 µL of 1 mg/mL ^13^C_10_ ATP; 10 µL of 1 mg/mL ^13^C ^15^N AMP; 10 µL of 1 mg/mL ^15^N ADP; and 20 µL of 100 µg/mL citric acid D_4_. For reduced buffer volumes (e.g. 25 ml), the internal standards were pre-diluted 1:2 (v:v) in extraction buffer and the same volumes were added as described above. The prepared extraction buffer was used immediately or stored at –20 °C for no more than one week to prevent the degradation of the internal standards. The culture medium was aspirated from each well, followed by two sequential washes with 1 mL of pre-warmed (37 °C) wash buffer to prevent thermal shock to the cells. If metabolite extraction was not performed immediately, the plates were snap-frozen in liquid nitrogen and stored at –80 °C. In parallel, two method blanks (wells without cells that were processed identically) were prepared to control for background metabolic contamination.

Metabolite extraction was performed by adding 400 µL of pre-cooled (−20 °C) extraction buffer to each well of the plate, which was placed on ice. The cells were then scraped thoroughly and transferred to pre-labelled 1.5 mL Eppendorf tubes, which were kept on ice. This extraction step was repeated twice more with fresh 400 µL aliquots of extraction buffer per well to ensure complete recovery of cellular material and precipitated proteins. All extracts from the same well were pooled into a single tube. These pooled extracts were then incubated for 40 minutes at 4 °C on a thermomixer, after which they were centrifuged at 21,000 × g for 10 minutes at 4 °C.

The resulting supernatant was transferred to new, pre-labelled 1.5 mL tubes. If required for multiple analyses, the extracts were divided into smaller portions. The remaining pellet was gently rinsed with a minimal volume of extraction buffer without disturbing it and stored at −80 °C for subsequent protein quantification or proteomic analyses.

The supernatant was completely dried in a SpeedVac concentrator set to 20 °C and 1000 rpm for 4–6 hours, depending on the sample volume. The dried metabolite pellets were then stored at –80 °C until further analysis.

### Metabolite extraction for lipid measurements

Metabolites were extracted from adherent cells cultured in 6-well plates, using a biphasic methyl tert-butyl ether (MTBE)-based protocol optimized for simultaneous lipid, polar metabolite, and protein recovery. Cell samples (∼1 × 10^6^ cells) were processed in biological replicates, alongside method blanks lacking cells to assess background contamination. The same wash buffer was used as before, and an extraction buffer prepared, consisting of 60% methanol (v/v, Optima LC-MS grade) in LCMS Ultra grade water, supplemented with internal standards: 10 µL of 2.5 mM U-^13^C^15^N-labelled amino acids; 10 µL of 1 mg/mL ^13^C_10_ ATP; 10 µL of 1 mg/mL ^13^C ^15^N_5_ AMP; 10 µL of 1 mg/mL ^15^N_5_ ADP; and 20 µL of 100 µg/mL citric acid D_4_ per 100 mL buffer. In addition, a lipid extraction buffer was prepared, composed of 50 mL MTBE (Sigma, 306975) with 20 µL EquiSPLASH Lipidomix (Avanti, 30731). Buffers were prepared fresh or stored up to one week at –20°C.

Cells were washed twice with 1 mL of pre-warmed wash buffer (37°C) and immediately processed or snap frozen in liquid nitrogen and stored at –80°C. For extraction, 400 µL of pre-cooled (–20°C) initial extraction buffer was added to each well, incubated for 10 min at –20°C, and cells were scraped and transferred into 2 mL tubes containing 900 µL of cold lipid extraction buffer. A second 400 µL wash of extraction buffer was added to the wells and pooled into the same tube. Tubes were mixed at 4°C and 1500 rpm for 30 min, then centrifuged at 21,000 × g for 10 min at 4°C.

The supernatant was transferred to a new tube and mixed with 200 µL LCMS Ultra grade water. After a second incubation at 15°C and 1500 rpm for 10 min, phase separation was achieved via centrifugation at 16,000 × g for 5 min. The upper (lipid-rich) phase (∼700 µL) was collected, aliquoted, and stored at – 80°C. The lower (polar metabolite) phase was cleaned of residual lipid, optionally aliquoted, dried in a SpeedVac concentrator at 20°C, and stored at –80°C. The protein pellet was washed and stored for proteomic analysis.

All steps involving MTBE were conducted under a fume hood. Samples were processed on ice or at low temperatures to minimize metabolic degradation.

### Anion-Exchange Chromatography Mass Spectrometry (AEX-MS) for the analysis of anionic metabolites

Extracted metabolites were re-suspended in 500 µl of UPLC/MS grade water (Biosolve), of which 100 µl were transferred to polypropylene autosampler vials (Chromatography Accessories Trott, Germany) before AEX-MS analysis.

The samples were analysed using a Dionex ionchromatography system (Integrion Thermo Fisher Scientific) as described previously^54^. In brief, 5 µL of the resuspended polar metabolite extract were injected in push-partial mode, using an overfill factor of 1, onto a Dionex IonPac AS11-HC column (2 mm × 250 mm, 4 μm particle size, Thermo Fisher Scientific) equipped with a Dionex IonPac AG11-HC guard column (2 mm × 50 mm, 4 μm, Thermo Fisher Scientific). The column temperature was held at 30°C, while the auto sampler temperature was set to 6°C. A potassium hydroxide gradient was generated using a potassium hydroxide cartridge (Eluent Generator, Thermo Scientific), which was supplied with deionized water (Milli-Q IQ 7000, Millipore). The metabolite separation was carried at a flow rate of 380 µL/min, applying the following gradient conditions: 0-3 min, 10 mM KOH; 3-12 min, 10−50 mM KOH; 12-19 min, 50-100 mM KOH; 19-22 min, 100 mM KOH, 22-23 min, 100-10 mM KOH. The column was re-equilibrated at 10 mM for 3 min.

For the analysis of metabolic pool sizes the eluting compounds were detected in negative ion mode using full scan measurements in the mass range m/z 77 – 770 on a Q-Exactive HF high resolution MS (Thermo Fisher Scientific). The heated electrospray ionization (ESI) source settings of the mass spectrometer were: Spray voltage 3.2 kV, capillary temperature was set to 300°C, sheath gas flow 50 AU, aux gas flow 14 AU at a temperature of 380°C and a sweep gas glow of 3 AU. The S-lens was set to a value of 40.

The LC-MS data analysis was performed using the TraceFinder software (Version 5.1, Thermo Fisher Scientific). The identity of each compound was validated by authentic reference compounds, which were measured at the beginning and the end of the sequence. For data analysis the area of the deprotonated [M-H^+^]^-1^ or doubly deprotonated [M-2H]^-2^ isotopologues mass peaks of every required compound were extracted and integrated using a mass accuracy <3 ppm and a retention time (RT) tolerance of <0.05 min as compared to the independently measured reference compounds. If no independent ^12^C experiments were carried out, where the pool size is determined from the obtained peak area of the ^12^C monoisotopologue, the pool size determination was carried out by summing up the peak areas of all detectable isotopologues per compound. These areas were then normalized, as performed for un-traced ^12^C experiments, to the internal standards, which were added to the extraction buffer, followed by a normalization to the protein content or the cell number of the analyzed samples. The relative isotope distribution of each isotopologue was calculated from the proportion of the peak area of each isotopologue towards the sum of all detectable isotopologues. The ^13^C enrichment, namely the area attributed to ^13^C molecules traced in the detected isotopologues, was calculated by multiplying the peak area of each isotopologue with the proportion of the ^13^C and the ^12^C carbon number for the corresponding isotopologue (the ^12^C and ^13^C monoisotopologue areas were multiplied with 0 and 1 respectively). The obtained ^13^C area of each isotopologue are summed up, providing the peak area fraction associated to ^13^C atoms in the compound. Dividing this absolute ^13^C area by the summed area of all isotopologues provides the relative ^13^C enrichment factor.

### Semi-targeted liquid chromatography-high-resolution mass spectrometry-based (LC-HRS-MS) analysis of amine-containing metabolites

The LC-HRMS analysis of amine-containing compounds was performed as described previously^55^. In brief: 50 µL of the available 500 µL of the above mentioned (AEX-MS) polar phase were mixed with 25 µl of 100 mM sodium carbonate (Sigma), followed by the addition of 25 µl 2% [v/v] benzoylchloride (Sigma) in acetonitrile (UPC/MS-grade, Biosove, Valkenswaard, Netherlands). Derivatized samples were thoroughly mixed and kept at 20°C until analysis.

For the LC-HRMS analysis, 1 µl of the derivatized sample was injected onto a 100 x 2.1 mm HSS T3 UPLC column (Waters). The flow rate was set to 400 µl/min using a binary buffer system consisting of buffer A (10 mM ammonium formate (Sigma), 0.15% [v/v] formic acid (Sigma) in UPC-MS-grade water (Biosove, Valkenswaard, Netherlands). Buffer B consisted of acetonitrile (IPC-MS grade, Biosove, Valkenswaard, Netherlands). The column temperature was set to 40°C, while the LC gradient was: 0% B at 0 min, 0-15% B 0- 4.1min; 15-17% B 4.1 – 4.5 min; 17-55% B 4.5-11 min; 55-70% B 11 – 11.5 min, 70-100% B 11.5 - 13 min; B 100% 13 - 14 min; 100-0% B 14 −14.1 min; 0% B 14.1-19 min; 0% B. The mass spectrometer (Q-Exactive Plus, Thermo Fisher Scientific) was operating in positive ionization mode recording the mass range m/z 100-1000. The heated ESI source settings of the mass spectrometer were: Spray voltage 3.5 kV, capillary temperature 300°C, sheath gas flow 60 AU, aux gas flow 20 AU at 330°C and the sweep gas was set to 2 AU. The RF-lens was set to a value of 60.

Semi-targeted data analysis for the samples was performed using the TraceFinder software (Version 4.1, Thermo Fisher Scientific). The identity of each compound was validated by authentic reference compounds, which were run before and after every sequence. Peak areas of [M + nBz + H]^+^ ions were extracted using a mass accuracy (<5 ppm) and a retention time tolerance of <0.05 min. Areas of the cellular pool sizes were calculated as described in the AEX-MS method.

### Isotope tracing experiment and mass spectrometry

Mouse embryonic stem cells were plated in a 6-well, grown for 24 h and transferred to media containing 100% labelled D-Glucose (U-^13^C_6_) (Sigma Aldrich) or supplemented with 10 mM ^13^C_2_-GlcNAc. The cells were exposed to the labelling medium (for 0, 30, 60, 360 min) before trypsinization and collection by centrifugation at 350 g (4 C°) for 3 minutes. The cells were washed in ice-cold PBS and the centrifugation step was repeated. The PBS was removed, and the cell pellet was frozen at −80°C until further procedure. On the day of the extraction, all tubes were placed on ice, and the samples were subjected to the respective extraction methods.

### 13C2 enrichment analysis of acetate after 3-Nitrophenolhydrazin (3-NPH) derivatization

Metabolites from frozen cell pellets were extracted using 150 µL of 80% methanol, containing 500 ng/mL of sodium acetate ^2^H_3_ (Sigma Aldrich, 176079). These samples were sonicated with 10 pulses of 0.5 seconds using a tip sonicator (Bandelin), to disrupt the frozen cell pellet within the extraction buffer. After sonication the extracts were incubated on a Thermomixer (Eppendorf) at 4°C and 1250 rpm. Extracts were cleared by centrifugation (15 min 15000 × g) and 30 µL of the cleared supernatant were used for derivatization.

The derivatization was performed based on the method described before ^56^ by adding and mixing sequentially 10 µL of 200 mM 3-NPH (Sigma, N21804), dissolved in 75% methanol and 10 µL of 96 mM N-(3-Dimethylaminopropyl)-N-ethyl-carbodiimide (EDC) dissolved methanol containing 6% pyridine within the cleared metabolite extract. The samples are mixed on a thermomixer (1250 rpm) at 30°C for 60 min, before adding 400 µL of ice-cold 50% methanol congaing 0.05% butylated hydroxytoluene (Sigma, W218405). Samples were subsequently incubated for 20 min at −20°C before clearing the supernatant with a 15min centrifugation at 21,000 × g at 4°C.

### Ultra-Performance Liquid Chromatography (UPLC) High Resolution Mass spectrometric (HRMS) analysis of derivatized ^13^C_2_ acetate

The UPLC-HRMS analysis was performed as described before ^57^, using a QE-Plus high-resolution mass spectrometer coupled to a Vanquish UHPLC chromatography system (Thermo Fisher Scientific). For the UPLC-HRMS analysis, 3 µl of the derivatized sample was injected onto a 100 × 2.1 mm HSS T3 UPLC column (Waters). The flow rate was set to 400 µl/min using a binary buffer system consisting of buffer A containing UPLC-grade water with 0.1% formic acid (Biosove, Valkenswaard, Netherlands). Buffer B consisted of acetonitrile:isopropanol 3:1 [v:v] containing 0.1% formic acid (Biosove, Valkenswaard, Netherlands). The column temperature was set to 40°C, while the LC gradient was: 3% B at 0 min, 3-100% B 0-9 min; 100% B 9-14 min; 100-3% B 14-14.5 min; 3% B 14.5-17 min. The mass spectrometer (Q-Exactive Plus) was operating in negative ionization mode recording the mass range m/z 100-1000. The heated ESI source settings of the mass spectrometer were: Spray voltage 3.0 kV, capillary temperature 300°C, sheath gas flow 50 AU, aux gas flow 10 AU at 330°C and the sweep gas was set to 1 AU. The RF-lens was set to a value of 40.

The UPLC-MS data analysis was performed using the TraceFinder software (Version 5.1, Thermo Fisher Scientific). For data analysis the area of the deprotonated acetate peak (C_2_H_4_O_2_) plus the (C_6_H_5_N_3_O) derivative of the 3-NPH (m/z 194.05710). Beyond to the monoisotopic acetate-3-NPH peak the M+1 and the M+2 peaks and the internal standard (acetate ^2^H_3_) were extracted and analyzed. The relative and absolute isotope distribution of each isotopologue was calculated from the proportion of the peak area of each isotopologue towards the sum of all detectable isotopologues. The ^13^C enrichment, namely the area attributed to ^13^C molecules traced in the detected isotopologues, was calculated by multiplying the peak area of each isotopologue with the proportion of the ^13^C and the ^12^C carbon number for the corresponding isotopologue (the ^12^C and ^13^C monoisotopologue areas were multiplied with 0 and 1 respectively). The obtained ^13^C area of each isotopologue are summed up, providing the peak area fraction associated to ^13^C atoms in the compound. Dividing this absolute ^13^C area by the summed area of all isotopologues provides the relative ^13^C enrichment factor.

### SDS-PAGE and Western blot analysis

Protein concentration of cell lysates was determined using the BCA protein assay kit (Pierce) according to the manufacturer’s instructions (Thermo Fisher Scientific). Samples were adjusted in 4x Laemmli sample buffer (BioRad) containing 50 mM β-mercaptoethanol. After boiling and a sonication step, equal protein amounts were subjected to SDS-PAGE and blotted on a nitrocellulose membrane using the Trans-Blot Turbo Transfer system (BioRad). All antibodies were used in 5% low-fat milk in TBS-Tween. After incubation with HRP-conjugated secondary antibody, the blot was developed using ECL solution (Merck Millipore) on a ChemiDoc MP Imaging System (BioRad).

The following antibodies were used in this study: SOX2 (ABclonal Cat# A19118, RRID:AB_2862611, 1:40,000), STAT3 (Cell Signaling Technology Cat# 9139, RRID:AB_331757, 1:1000), NANOG (Proteintech Cat# 14295-1-AP, RRID:AB_1607719; 1:100), GFPT2 (Abcam Cat# ab190966, RRID:AB_2868470, 1:5000), GNPNAT1 (ABclonal Cat# A17759, RRID:AB_2769647, 1:1000), FLAG (Sigma-Aldrich Cat# F1804, RRID:AB_262044, 1:2000), AMDHD2 (ms, S6 clone, produced by Bernhard Schermer, 1:500), FASN (ABclonal Cat# A21182, 1:1000), ACSS2/AceCS1 (Cell Signaling Technology Cat# 3658, RRID:AB_2222710, 1:1000), ACLY (Proteintech Cat# 15421-1-AP, RRID:AB_2223741, 1:1000), GFP (BioLegend Cat# 902602, RRID:AB_2565022, 1:5000), HSP90 (Cell Signaling Technology Cat# 4874S, RRID:AB_2121214, 1:5000), alpha-TUBULIN (Sigma-Aldrich Cat# T6074, RRID:AB_477582, 1:15000), V5 (Sigma-Aldrich Cat# V8012, RRID:AB_261888 1:2000), beta-ACTIN (Abcam Cat# ab8224, RRID:AB_449644, 1:1000), rabbit IgG (Thermo Fisher Scientific Cat# G-21234, RRID:AB_2536530,1:5000), and mouse IgG (Thermo Fisher Scientific Cat# G-21040, RRID:AB_2536527, 1:5000).

### Mass spectrometry-based proteomics

Cells were lysed by adding lysis buffer (6 M Guanidinium chloride, 2.5 mM Tris(2-carboxyethyl)phosphine, 10 mM chloroacetamide, 100 mM Tris-HCL) and heating up to 95°C for 10 min at 700 rpm. For complete denaturation of proteins, samples were sonicated for 10 min (30 s on/30 s off). Afterwards, samples were centrifuged at 20.000 g for 20 min at room temperature and supernatants were transferred to new tubes. 100 µg of protein were digested with trypsin (1:200 w/w, Promega) over night at 37 °C and 800 rpm. The next day the digest was stopped by acidification with formic acid and peptides were purified by Pierce Peptide Desalting Spin columns (Thermo Fisher Scientific) according to the manufacturer’s protocol. Two micrograms of desalted peptides were dried out and reconstituted.

The Evosep One liquid chromatography system was used for analyzing the samples with the predefined 30 samples per day (15SPD or 30SPD) method ^58^. The analytical column we used was an ReproSil-Pur column, 15 cm × 150 µm, with 1.9 µm C18 beads (EV1106 Endurance Column, Evosep). The mobile phases A and B were 0.1 % formic acid in water and 0.1% formic acid in 100% ACN, respectively. Peptides were analyzed on a hybrid TIMS quadrupole TOF mass spectrometer (timsTOF Pro 2 or timsTOF HT, Bruker) in a data-independent acquisition parallel accumulation, serial fragmentation (diaPASEF) mode. The mass spectra range was set to 100 - 1700 m/z and TIMS ion accumulation and ramp times were set to 100 ms and total cycle time was 2.0 s. The ion mobility range was set to 1/K0 = 0.8 - 1.25 V-s/cm2. Isolation windows in the m/z versus ion mobility plane were defined to cover the region of highest precursor ion density with an m/z slice width of 26 Th. Collision energy was applied linearly with ion mobility from 0.6 to 2.0 V-s/cm2, and collision energy from 20 to 59 eV. Raw data was analyzed using Spectronaut version 17.4, 18.0, or 19.3 (Biognosys) using the default parameters against the one-protein-per-gene reference proteome for Mus musculus, UP000000589, downloaded August, 2022. Methionine oxidation and protein N-terminal acetylation were set as variable modifications; cysteine carbamidomethylation was set as fixed modification. The digestion parameters were set to “specific” and “Trypsin/P,” with two missed cleavages permitted. Protein groups were filtered for at least two valid values in at least one comparison group and missing values were imputed from a normal distribution with a down-shift of 1.8 and standard deviation of 0.3. Differential expression analysis was performed using limma, version 3.60.6, in R, version 4.4.0^59^. Gene set enrichment analysis (GSEA) was performed using WebGestaltR version 0.4.6^60^. Clustering and visualization of the GSEA results was done using aPEAR version 1.0.0^61^.

### CUT&RUN for H3ac, H3K27ac and H3K27me3

CUT&RUN (Cleavage Under Targets and Release Using Nuclease) was performed for WT and AMDHD2 KO AN3-12 lines following the EpiCypher CUT&RUN protocol with some modifications (v2.0). A total of 500,000 cells were harvested and washed with wash buffer (as described in the protocol with the addition of 5 mM sodium butyrate), then combined with 50,000 *Drosophila melanogaster* nuclei serving as spike-in controls. The pooled cell-nuclei mixture was subsequently incubated with activated Concanavalin A-coated magnetic beads (BioMagPlus Concanavalin A, #BP531) for immobilization. Antibody incubation was performed overnight at 4°C with gentle nutation using H3ac (Active Motif Cat# 39139, RRID:AB_2687871, 1:50 dilution), H3K27ac (Active Motif Cat# 39133, RRID:AB_2561016, 1:50 dilution), H3K27me3 (Active Motif Cat# 39155, RRID:AB_2561020, 1:50 dilution), and Normal Rabbit IgG control (Cell Signaling Technology Cat# 2729, RRID:AB_1031062, 1:50 dilution) antibodies. Samples were incubated with protein AG-MNase (Epicypher, #15-1016) fusion protein for 2 hours at 4°C. Sequencing libraries were prepared using the EpiCypher CUT&RUN Library Prep kit according to manufacturer’s protocol. DNA samples were purified and concentrated using the Zymo DNA Clean & Concentrator Kit, followed by elution in the kit-supplied buffer. Final library concentrations were quantified using the Qubit dsDNA High Sensitivity Assay (Thermo Fisher Scientific), and library quality and fragment size distribution were assessed using the High Sensitivity DNA Kit on an Agilent 2100 Bioanalyzer (Agilent Technologies). Libraries were sequenced on an Illumina NextSeq 2000 platform to generate 59 bp paired-end reads.

### CUT&RUN data analysis

Raw sequencing data was processed and analyzed using the nf-core/cutandrun pipeline (v3.2.2)^62^, which includes quality control, adapter trimming, alignment to the reference genome (mm10 for target reads and BDGP6 for spike-in reads), spike-in normalization and peak calling. Peak-calling was done using MACS2 (v2.2.7.1) with broad and narrow peak calling, as appropriate. Differential peak analysis was performed utilizing Diffbind (v3.4.11), GO enrichment was performed with clusterProfiler (v4.16.0) and deepTools (v3.5.1) was used to compute global signal distribution.

### Statistical analysis

The mean of technical replicates is plotted for each biological replicate. Biological replicates represent different passages of the cells that were seeded on independent days. Statistical significance was calculated using GraphPad Prism (GraphPad Software, San Diego, California). The statistical test used is indicated in the respective figure legend. Significance levels are * p<0.05, ** p<0.01, *** p<0.001 relative to the respective control.

## Data availability

All raw and analysed data were deposited on Mendeley Data. The CUT&RUN sequencing data were deposited on the Gene Expression Omnibus platform.

---

**Supplementary Table 1.**
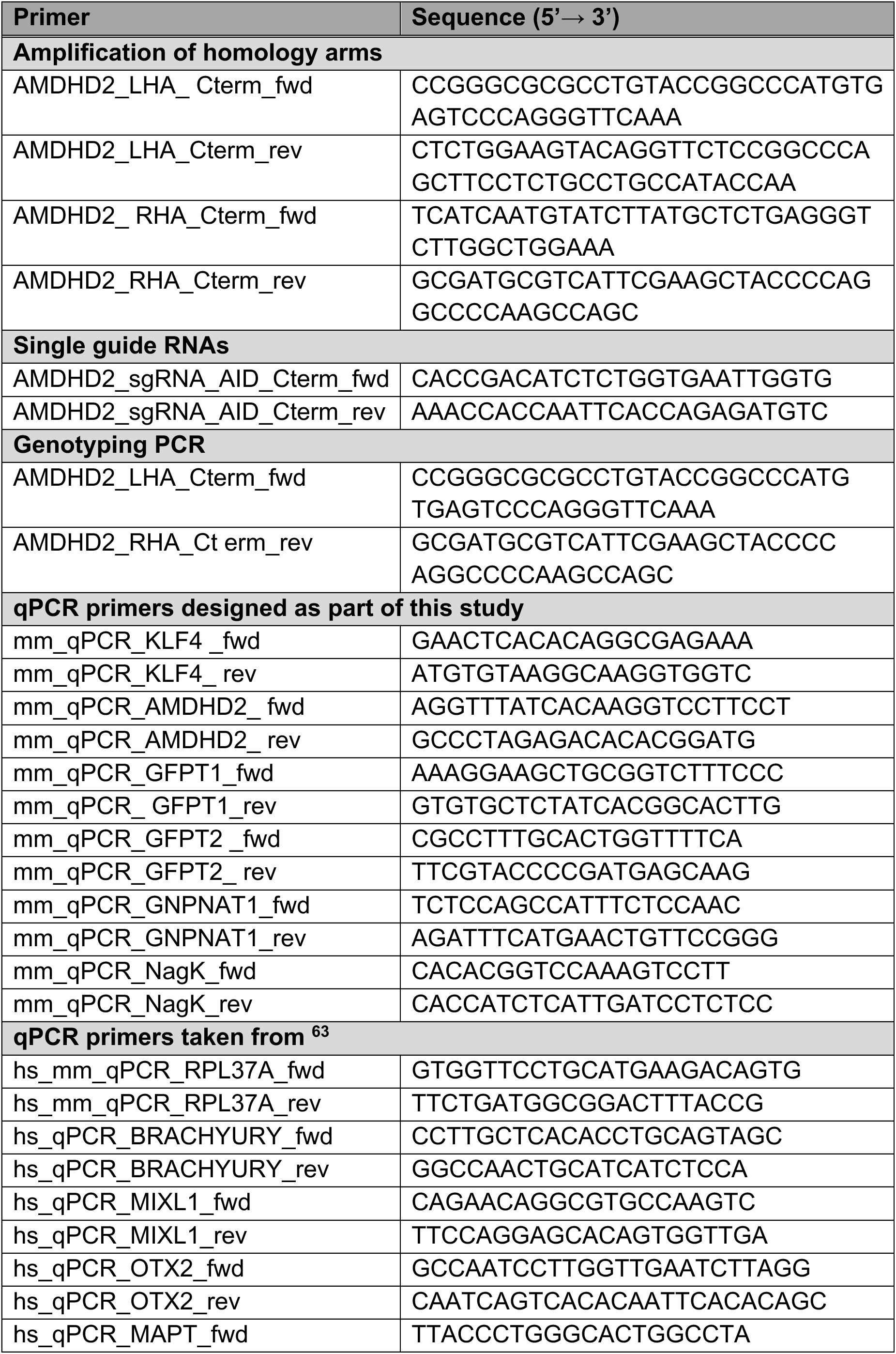
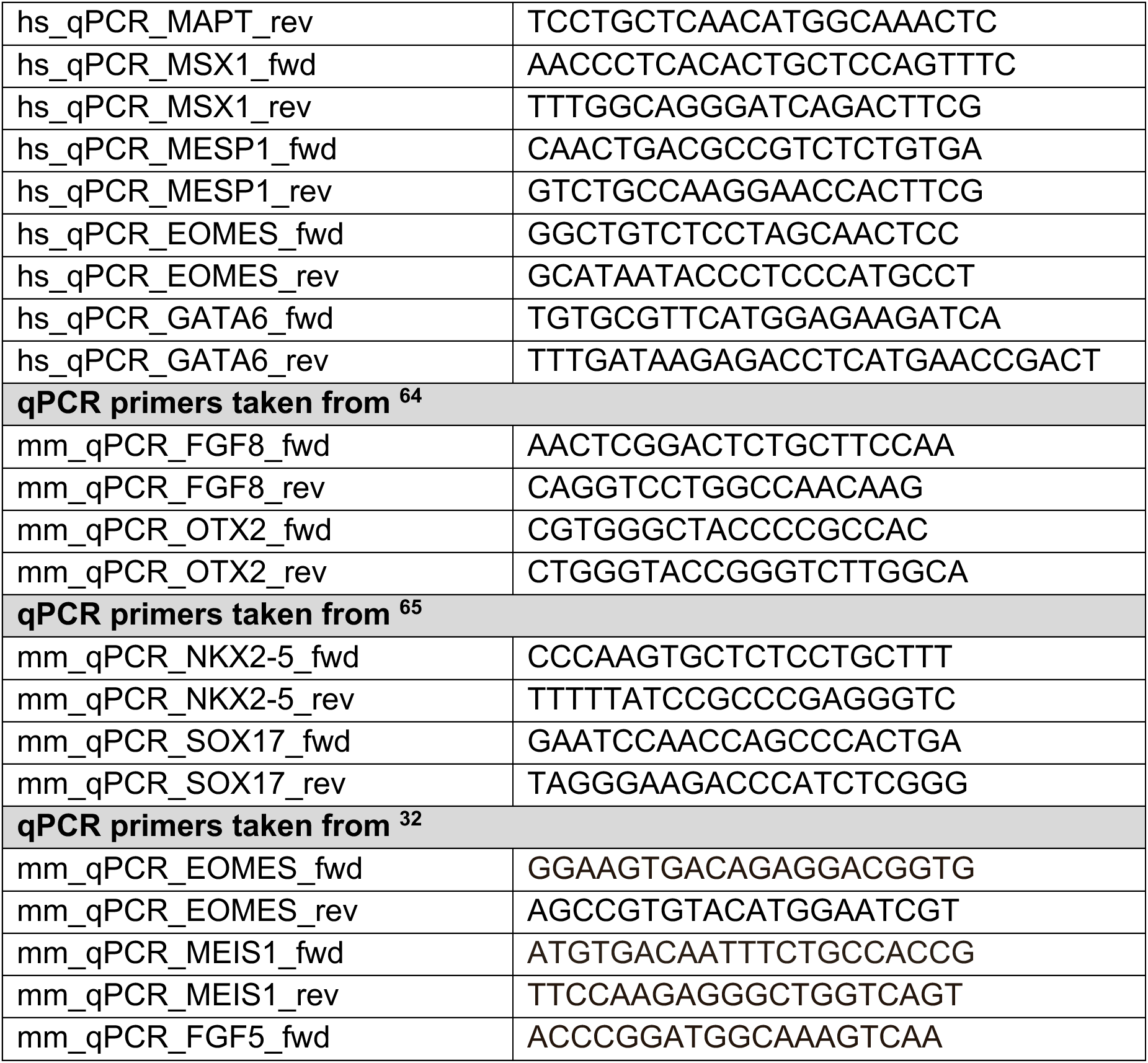
Primer and sgRNA sequences used in this study.

## Acknowledgments

The C-terminal degron tagging plasmid was kindly provided by F. Dossin (Institut Pasteur). We thank F. Dethloff, and Y. Hinze from the MPI-AGE metabolomics core facility. Moreover, we thank I. Atanassov, T. Colby, X. Li and I. Matic from the MPI-AGE proteomics core facility. We thank L. Schumacher, M. Germer and C. Kukat from the MPI-AGE FACS and imaging core facility. We thank A.Iqbal, M. Duhamel and J. Boucas from the MPI-AGE Bioinformatics facility. Finally, the UKBi017-A iPSC line was provided by Michael Peitz and Oliver Brüstle, Cell Programming Core Facility and Institute of Reconstructive Neurobiology, Medical Faculty of the University of Bonn. This work was supported by the German Research Foundation (DFG, Project no. DE 2326/3-1) and by the Max Planck Society. L.H. was supported by the Jürgen Manchot Foundation.

## Author contributions

V.K. contributed to the conceptualisation, investigation and supervision of the study, as well as the writing of the original manuscript. L.H. executed all qPCR analyses, generated the AMDHD2-AID AN3-12 mESC line and also prepared numerous metabolic measurements with the help of J.F. Y.B.A. performed all CUT&RUN experiments under the supervision of P.T., while A.H. and S.M.A. generated the PancACe sensor AN3-12 mESC line under the supervision of N.T., who also performed all flow cytometry experiments. L.W. performed the iPSC differentiation experiments under the supervision of H.S. P.G. designed and executed all metabolic measurements. M.S.D. initiated the study and obtained funding. I.H. was responsible for project administration, conceptualisation, resources, and supervision of the study, as well as the writing of the original manuscript.

## References

1. Folmes, C.D. et al. Somatic oxidative bioenergetics transitions into pluripotency-dependent glycolysis to facilitate nuclear reprogramming. Cell metabolism 14, 264–271 (2011).

2. Panopoulos, A.D. et al. The metabolome of induced pluripotent stem cells reveals metabolic changes occurring in somatic cell reprogramming. Cell research 22, 168–177 (2012).

3. Yanes, O. et al. Metabolic oxidation regulates embryonic stem cell differentiation. Nature chemical biology 6, 411–417 (2010).

4. Kim, H. et al. Core Pluripotency Factors Directly Regulate Metabolism in Embryonic Stem Cell to Maintain Pluripotency. Stem Cells 33, 2699–2711 (2015).

5. Hsieh, M.-H. et al. p63 and SOX2 Dictate Glucose Reliance and Metabolic Vulnerabilities in Squamous Cell Carcinomas. Cell Reports 28, 1860–1878.e1869 (2019).

6. Chen, C.-L. et al. NANOG Metabolically Reprograms Tumor-Initiating Stem-like Cells through Tumorigenic Changes in Oxidative Phosphorylation and Fatty Acid Metabolism. Cell Metabolism 23, 206–219 (2016).

7. Moussaieff, A. et al. Glycolysis-mediated changes in acetyl-CoA and histone acetylation control the early differentiation of embryonic stem cells. Cell Metabolism 21, 392–402 (2015).

8. Kuna, R.S. et al. Inter-organelle cross-talk supports acetyl-coenzyme A homeostasis and lipogenesis under metabolic stress. Science Advances 9, eadf0138 (2023).

9. Li, C., Liu, W., Liu, Y., Wang, W. & Deng, W. Role of ATP citrate lyase and its complementary partner on fatty acid synthesis in gastric cancer. Scientific reports 14, 30043 (2024).

10. Zhao, S. et al. ATP-Citrate Lyase Controls a Glucose-to-Acetate Metabolic Switch. Cell Reports 17, 1037–1052 (2016).

11. Zaidi, N., Royaux, I., Swinnen, J.V. & Smans, K. ATP Citrate Lyase Knockdown Induces Growth Arrest and Apoptosis through Different Cell- and Environment-Dependent Mechanisms. Molecular Cancer Therapeutics 11, 1925–1935 (2012).

12. Marshall, S., Bacote, V. & Traxinger, R.R. Discovery of a metabolic pathway mediating glucose-induced desensitization of the glucose transport system. Role of hexosamine biosynthesis in the induction of insulin resistance. J Biol Chem 266, 4706–4712 (1991).

13. Ghosh, S., Blumenthal, H.J., Davidson, E. & Roseman, S. Glucosamine metabolism. V. Enzymatic synthesis of glucosamine 6-phosphate. J Biol Chem 235, 1265–1273 (1960).

14. Wellen, K.E. et al. The hexosamine biosynthetic pathway couples growth factor-induced glutamine uptake to glucose metabolism. Genes Dev 24, 2784–2799 (2010).

15. Wang, J., Liu, X., Liang, Y.H., Li, L.F. & Su, X.D. Acceptor substrate binding revealed by crystal structure of human glucosamine-6-phosphate N-acetyltransferase 1. FEBS Lett 582, 2973–2978 (2008).

16. Arreola, R., Valderrama, B., Morante, M.L. & Horjales, E. Two mammalian glucosamine-6-phosphate deaminases: a structural and genetic study. FEBS Lett 551, 63–70 (2003).

17. Weidanz, J.A. et al. N-acetylglucosamine kinase and N-acetylglucosamine 6-phosphate deacetylase in normal human erythrocytes and Plasmodium falciparum. British journal of haematology 95, 645–653 (1996).

18. Bergfeld, A.K., Pearce, O.M., Diaz, S.L., Pham, T. & Varki, A. Metabolism of vertebrate amino sugars with N-glycolyl groups: elucidating the intracellular fate of the non-human sialic acid N-glycolylneuraminic acid. J Biol Chem 287, 28865–28881 (2012).

19. Ricciardiello, F. et al. Inhibition of the Hexosamine Biosynthetic Pathway by targeting PGM3 causes breast cancer growth arrest and apoptosis. Cell Death Dis 9, 377 (2018).

20. Mio, T., Yabe, T., Arisawa, M. & Yamada-Okabe, H. The eukaryotic UDP-N-acetylglucosamine pyrophosphorylases. Gene cloning, protein expression, and catalytic mechanism. J Biol Chem 273, 14392–14397 (1998).

21. Chiaradonna, F., Ricciardiello, F. & Palorini, R. The Nutrient-Sensing Hexosamine Biosynthetic Pathway as the Hub of Cancer Metabolic Rewiring. Cells 7 (2018).

22. Gloster, T.M. et al. Hijacking a biosynthetic pathway yields a glycosyltransferase inhibitor within cells. Nature chemical biology 7, 174–181 (2011).

23. Jang, H. et al. O-GlcNAc regulates pluripotency and reprogramming by directly acting on core components of the pluripotency network. Cell Stem Cell 11, 62–74 (2012).

24. Myers, S.A. et al. SOX2 O-GlcNAcylation alters its protein-protein interactions and genomic occupancy to modulate gene expression in pluripotent cells. Elife 5, e10647 (2016).

25. Constable, S., Lim, J.-M., Vaidyanathan, K. & Wells, L. O-GlcNAc transferase regulates transcriptional activity of human Oct4. Glycobiology 27, 927–937 (2017).

26. Omelková, M. et al. An O-GlcNAc transferase pathogenic variant linked to intellectual disability affects pluripotent stem cell self-renewal. Disease Models & Mechanisms 16, dmm049132 (2023).

27. Shi, F.-T. et al. Ten-Eleven Translocation 1 (Tet1) Is Regulated by O-Linked N-Acetylglucosamine Transferase (Ogt) for Target Gene Repression in Mouse Embryonic Stem Cells*. Journal of Biological Chemistry 288, 20776–20784 (2013).

28. Vella, P. et al. Tet proteins connect the O-linked N-acetylglucosamine transferase Ogt to chromatin in embryonic stem cells. Mol Cell 49, 645–656 (2013).

29. Parween, S. et al. Nutrient sensitive protein O-GlcNAcylation modulates the transcriptome through epigenetic mechanisms during embryonic neurogenesis. Life Science Alliance 5, e202201385 (2022).

30. Sakabe, K., Wang, Z. & Hart, G.W. Beta-N-acetylglucosamine (O-GlcNAc) is part of the histone code. Proc Natl Acad Sci U S A 107, 19915–19920 (2010).

31. Nishihara, S. Glycans in stem cell regulation: from Drosophila tissue stem cells to mammalian pluripotent stem cells. FEBS Letters 592, 3773–3790 (2018).

32. Kroef, V. et al. GFPT2/GFAT2 and AMDHD2 act in tandem to control the hexosamine pathway. Elife 11 (2022).

33. Vander Heiden, M.G., Cantley, L.C. & Thompson, C.B. Understanding the Warburg effect: the metabolic requirements of cell proliferation. Science 324, 1029–1033 (2009).

34. Manganelli, G. et al. Modulation of the pentose phosphate pathway induces endodermal differentiation in embryonic stem cells. PLoS One 7, e29321 (2012).

35. Hocke, G.M., Cui, M.Z. & Fey, G.H. The LIF response element of the alpha 2 macroglobulin gene confers LIF-induced transcriptional activation in embryonal stem cells. Cytokine 7, 491–502 (1995).

36. Folmes, Clifford D.L., Dzeja, Petras P., Nelson, Timothy J. & Terzic, A. Metabolic Plasticity in Stem Cell Homeostasis and Differentiation. Cell Stem Cell 11, 596–606 (2012).

37. Campbell, S. et al. Glutamine deprivation triggers NAGK-dependent hexosamine salvage. Elife 10, e62644 (2021).

38. Dossin, F. et al. SPEN integrates transcriptional and epigenetic control of X-inactivation. Nature 578, 455–460 (2020).

39. Pouikli, A. et al. Chromatin remodeling due to degradation of citrate carrier impairs osteogenesis of aged mesenchymal stem cells. Nature Aging 1, 810–825 (2021).

40. Smith, J.J. et al. A genetically encoded fluorescent biosensor for visualization of acetyl-CoA in live cells. Cell Chemical Biology 32, 325–337.e310 (2025).

41. Friedman, S. & Fraenkel, G. Reversible enzymatic acetylation of carnitine. Archives of biochemistry and biophysics 59, 491–501 (1955).

42. Meissen, J.K. et al. Induced pluripotent stem cells show metabolomic differences to embryonic stem cells in polyunsaturated phosphatidylcholines and primary metabolism. PLoS One 7, e46770 (2012).

43. Wang, L. et al. Fatty acid synthesis is critical for stem cell pluripotency via promoting mitochondrial fission. EMBO J 36, 1330–1347 (2017).

44. Yang, Y.R. et al. O-GlcNAcase is essential for embryonic development and maintenance of genomic stability. Aging cell 11, 439–448 (2012).

45. Marsboom, G. et al. Glutamine metabolism regulates the pluripotency transcription factor OCT4. Cell reports 16, 323–332 (2016).

46. Speakman, C.M. et al. Elevated O-GlcNAc levels activate epigenetically repressed genes and delay mouse ESC differentiation without affecting naïve to primed cell transition. Stem Cells 32, 2605–2615 (2014).

47. Gnoni, G.V., Priore, P., Geelen, M.J. & Siculella, L. The mitochondrial citrate carrier: metabolic role and regulation of its activity and expression. IUBMB life 61, 987–994 (2009).

48. Khacho, M. & Slack, R.S. Mitochondrial activity in the regulation of stem cell self-renewal and differentiation. Current Opinion in Cell Biology 49, 1–8 (2017).

49. Willnow, P. & Teleman, A.A. Nuclear position and local acetyl-CoA production regulate chromatin state. Nature 630, 466–474 (2024).

50. Moussaieff, A. et al. Glycolysis-mediated changes in acetyl-CoA and histone acetylation control the early differentiation of embryonic stem cells. Cell Metab 21, 392–402 (2015).

51. Ma, S. et al. Nutrient-driven histone code determines exhausted CD8+ T cell fates. Science 0, eadj3020 (2024).

52. Zhu, R. et al. ACSS2 acts as a lactyl-CoA synthetase and couples KAT2A to function as a lactyltransferase for histone lactylation and tumor immune evasion. Cell Metabolism (2024).

53. Huppertz, I. et al. Riboregulation of Enolase 1 activity controls glycolysis and embryonic stem cell differentiation. Molecular Cell 82, 2666–2680.e2611 (2022).

54. Schwaiger, M. et al. Anion-Exchange Chromatography Coupled to High-Resolution Mass Spectrometry: A Powerful Tool for Merging Targeted and Non-targeted Metabolomics. Anal Chem 89, 7667–7674 (2017).

55. Bonekamp, N.A. et al. Small-molecule inhibitors of human mitochondrial DNA transcription. Nature 588, 712–716 (2020).

56. Xie, G. et al. A Metabolite Array Technology for Precision Medicine. Anal Chem 93, 5709–5717 (2021).

57. Hodek, O., Henderson, J., Argemi-Muntadas, L., Khan, A. & Moritz, T. Structural elucidation of 3-nitrophenylhydrazine derivatives of tricarboxylic acid cycle acids and optimization of their fragmentation to boost sensitivity in liquid chromatography-mass spectrometry. J Chromatogr B Analyt Technol Biomed Life Sci 1222, 123719 (2023).

58. Bache, N. et al. A Novel LC System Embeds Analytes in Pre-formed Gradients for Rapid, Ultra-robust Proteomics*. Molecular & Cellular Proteomics 17, 2284–2296 (2018).

59. Ritchie, M.E. et al. limma powers differential expression analyses for RNA-sequencing and microarray studies. Nucleic Acids Res 43, e47 (2015).

60. Liao, Y., Wang, J., Jaehnig, E.J., Shi, Z. & Zhang, B. WebGestalt 2019: gene set analysis toolkit with revamped UIs and APIs. Nucleic Acids Res 47, W199–w205 (2019).

61. Kerseviciute, I. & Gordevicius, J. aPEAR: an R package for autonomous visualization of pathway enrichment networks. Bioinformatics 39 (2023).

62. Cheshire, C. et al. nf-core/cutandrun: nf-core/cutandrun v3.2.2 Iridium Ibis. Zenodo **v3.2.2** (2024).

63. Bartsch, D. et al. mRNA translational specialization by RBPMS presets the competence for cardiac commitment in hESCs. Sci Adv 9, eade1792 (2023).

64. Huang, X. et al. Zfp281 is essential for mouse epiblast maturation through transcriptional and epigenetic control of Nodal signaling. Elife 6 (2017).

65. Elling, U. et al. A reversible haploid mouse embryonic stem cell biobank resource for functional genomics. Nature 550, 114–118 (2017).

